# Transglutaminase 2 is an RNA-binding protein: Experimental verification and characterisation of a novel transglutaminase feature

**DOI:** 10.1101/2024.04.26.591323

**Authors:** Bianka Csaholczi, Anna Renáta Csuth, Ilma Rita Korponay-Szabó, László Fésüs, Róbert Király

**Affiliations:** Department of Biochemistry and Molecular Biology, Faculty of Medicine, University of Debrecen, Debrecen, Hungary; Doctoral School of Molecular Cell and Immune Biology, University of Debrecen, Debrecen, Hungary; Department of Pediatrics, Faculty of Medicine, University of Debrecen, Debrecen, Hungary

**Keywords:** transglutaminase 2, RNA-binding, RNA-binding protein, endothelial cells

## Abstract

Transglutaminase 2 (TG2) is a uniquely versatile protein with diverse catalytic activities, such as transglutaminase, protein disulfide isomerase, GTPase, protein kinase, and participates in several biological processes. According to information available in the RBP2GO database, TG2 can be an RNA-binding protein (RBP). RBPs participate in posttranscriptional gene expression regulation, influencing RNAs’ function, while RNA molecules can also modulate RBPs’ biological activity. Our goal was to confirm this novel character of TG2 in human umbilical cord vein endothelial cells (HUVEC), which physiologically express TG2.

First, UV cross-linked RNA-protein complexes were isolated from immortalised HUVEC using orthogonal organic phase separation. Compared with the RBP2GO database, mass spectrometry identified 392 potential RBPs, including TG2 and 20 novel, endothelium-related RBPs. Total RNA from HUVEC pulled down recombinant human TG2. Complex formation between TG2 and a 43-mer RNA molecule with a secondary structure as well as a homo-oligomeric single-stranded poly(dG), but not poly(dA), could be observed in magnetic RNA-protein pull-down experiments. Experiments with TG2 inhibitors NC9 and GTPγS, which stabilise its open and closed conformation, respectively, revealed that the open conformation of the enzyme favoured RNA-binding. Biolayer interferometry revealed a high binding affinity between TG2 and RNA with a K_D_ value of 88 nM.

We propose that superficial residues on the catalytic core and C-terminal β-barrel domains, being in a hidden position in the closed TG2, are involved in RNA binding. Our study demonstrates TG2’s previously uncharacterised RNA-binding ability, opening new avenues for understanding its multi-functionality.

**Graphical abstract:** 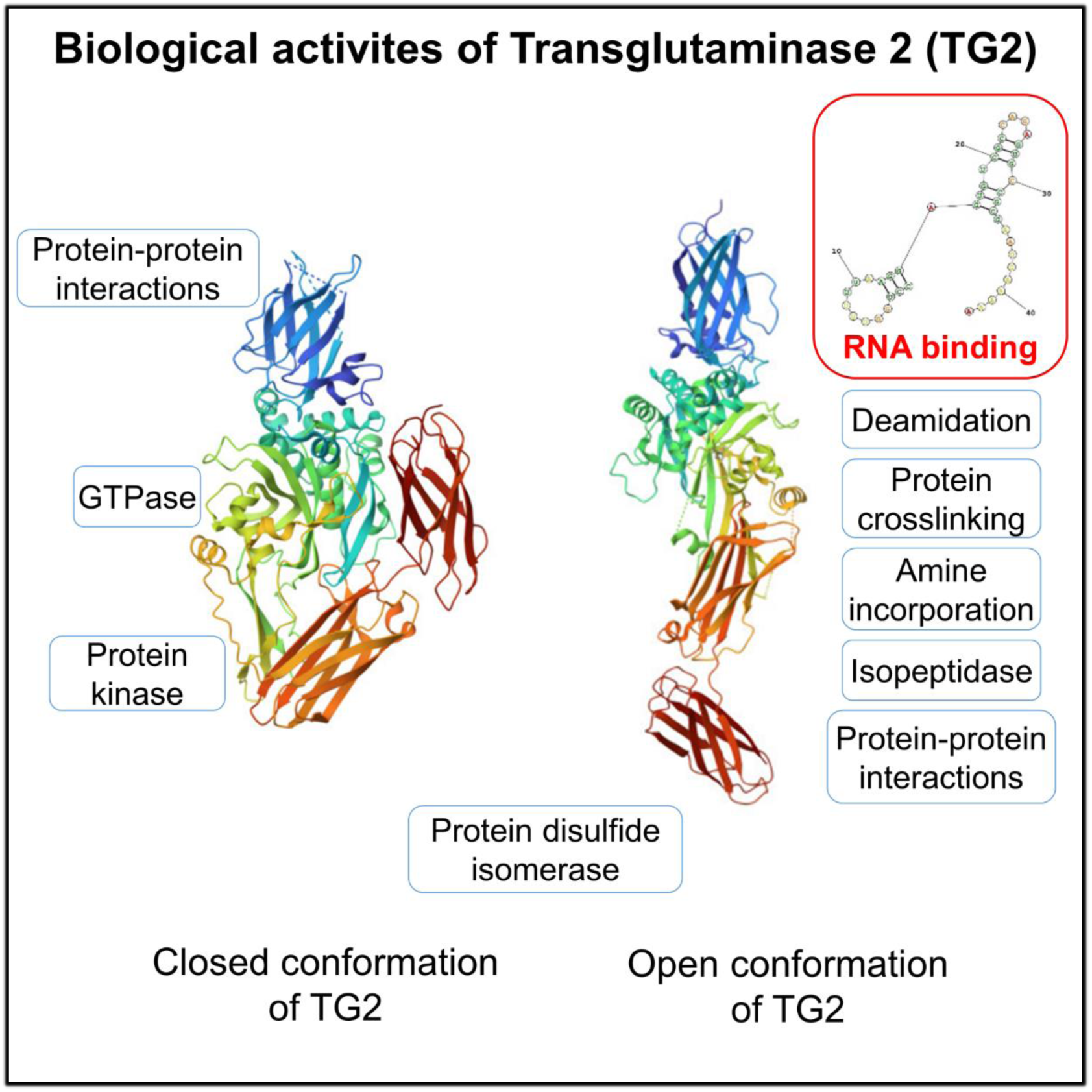

Transglutaminase 2 (TG2) is a unique multifunctional protein demonstrating large conformational changes. It has various transglutaminase and other catalytic and non-catalytic activities which show conformation dependency. Our study has characterised a novel, open conformation-related biological activity of TG2 and its RNA-binding ability.

## Introduction

Human transglutaminase 2 (TG2) is a uniquely versatile, ubiquitous protein in the human body. It has several diverse catalytic activities; as a Ca^2+^-dependent transglutaminase, it can crosslink glutamine and lysine residues of proteins or incorporate biogenic amines into protein glutamine side chains (transamidase activity), but it can revert the previously formed isopeptide bond by its isopeptidase activity or hydrolyse protein glutamine to glutamate residue (Fesus & Piacentini; 2002). In addition, it has GTPase, ATPase (Iismaa et al., 1997), protein disulfide isomerase (Hasegawa et al., 2003) and protein kinase activities (Mishra & Murphy, 2004). Besides its enzymatic activities, TG2 participates in various biological processes, such as cell adhesion, migration, angiogenesis, phagocytosis, regulation of mitochondrial processes and cell survival or death (Eckert et al., 2014). Its known multi-functionality implies the necessity of complex structure and regulatory features. TG2 is composed of an N-terminal β-sandwich, catalytic core and two C-terminal β-barrel domains (Demény et al., 2015). In the presence of GTP/GDP TG2 has a compacted (so-called closed) conformation, which turns to a relaxed open conformation in the presence of increasing Ca^2+^ concentrations. In this way, GTP and Ca^2+^ binding regulate reciprocally the GTPase and transglutaminase activities followed by a large conformational change (Király et al., 2011; Jeong et al., 2020). In addition, extracellular oxidative factors (Basso et al., 2012) or NO (Santhanam et al., 2010; Lai et al., 2017) in endothelial cells lead to the inhibition of transglutaminase activities via the formation of disulfide bonds or nitrosylation of Cys residues, respectively.

There are increasing pieces of evidence which show the involvement of TG2 in the regulation of gene expression. It can serotonylate the histone tails (Farrelly et al., 2019), affecting transcription. Silencing (Nadalutti et al., 2011) or knocking out TG2 (Yunes-Medina et al., 2018) results in changes in the expression of several genes. TG2 is involved in neurodegenerative diseases where transcriptional dysregulation plays a significant pathogenic role. (McConoughey et al., 2010). This previous information has led to our assumption that TG2 may bind RNA and, in this way, plays a role in posttranscriptional gene expression regulation. RNA-binding proteins (RBPs) can modulate the stability, processing and localisation of various RNA molecules and the translation of mRNAs (Hentze et al., 2018). Riboregulation occurs when RNA influences the stability, localisation, function, and activity of proteins (Guiducci et al., 2019). RBP2GO, a large and up-to-date RNA-binding protein database (Caudron-Herger et al., 2021), has a record which lists TG2 as one of the RBPs in the U2OS osteosarcoma cell line (Queiroz et al., 2019).

In our study, we aimed to confirm experimentally the RNA-binding ability of TG2 by using an immortalised human umbilical cord vein endothelial cell line (HUVEC; Shaw et al., 2021) that physiologically expresses TG2 in high amounts. We have identified RBPs from these cells using orthogonal organic phase separation (OOPS) and found TG2 among them. Subsequently, we demonstrated that desthiobiotin-labelled total RNA from the HUVEC can pull down recombinant TG2 from cell protein extract. Luckily, we have found a 43-mer RNA sequence which can pull down either HUVEC-produced endogenous or bacterially expressed recombinant human TG2. TG2-RNA interaction showed TG2 conformation, nucleic acid sequence and secondary structure dependence, and a middle nanomolar K_D_ value determined by a Blitz (biolayer interference) method. According to our knowledge, this is the first study which has confirmed the novel RNA-binding feature of TG2.

## Results

### Transglutaminase 2 is among the RNA-binding proteins of HUVEC

In order to confirm the potential RNA-binding property of TG2, we used an orthogonal organic phase separation (OOPS) method in an immortalised HUVEC line. OOPS is an economical and simple way to isolate RBPs from various cells. To generate a sufficient amount of cell samples for successful subsequent LC-MS/MS identification, we used a HUVEC line immortalised earlier by telomerase transduction (Shaw et al., 2021). This cell line maintained the expression of TG2 and classical endothelial cell markers analysed by mRNA sequencing (Supplementary Fig. 1).

During RBP isolation, the UV-crosslinked RNA-protein-containing interphase was washed three times until the last organic phase wash contained only very low amounts of TG2 according to Western blot (Fig. 1). In the case of UV-treated cells, after RNA degrading treatments (sonication, denaturation, RNase A digestion), the separated organic phase showed a strong TG2 signal on Western blot when the same solvent volumes were applied during the separation steps.

**Figure 1.**
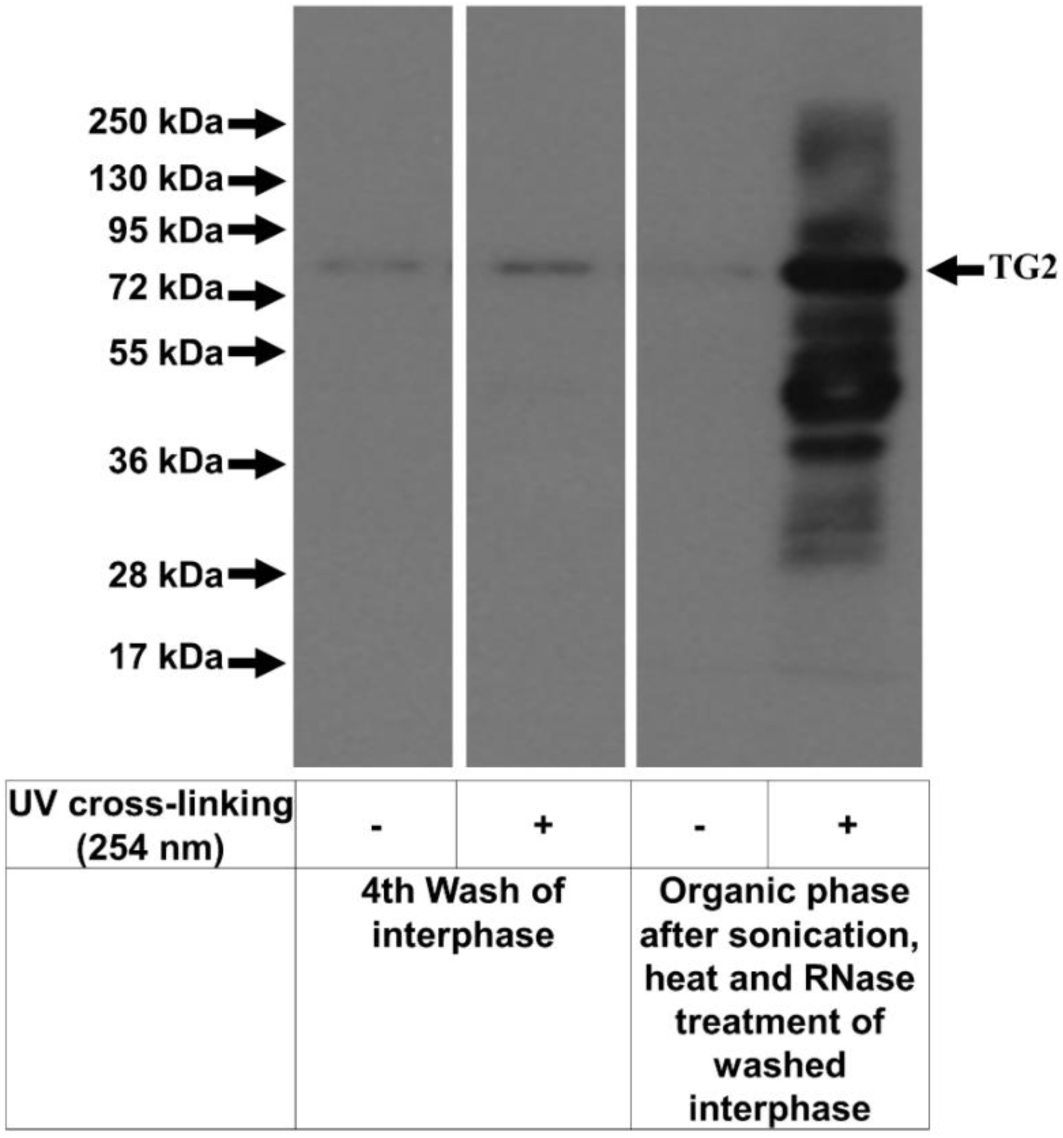
TG2 was identified among the RNA-binding proteins of the HUVEC line isolated using orthogonal organic phase separation. A representative Western blot showing the elimination of free TG2 from the interphase containing UV-crosslinked RNA-protein complexes and the appearance of TG2 in the organic phase upon RNA degradation. Equal volumes of the organic phases were precipitated and resolved in the same sample volume for Western blotting.

To investigate the relative abundance of TG2 among the RNA-binding proteins of HUVEC cells, the isolated RBP fractions were subjected to LC-MS/MS analysis. During the evaluation of the raw mass spectrometric data, each protein representing at least one unique tryptic peptide with four subsequent degradation ions was considered as a hit. Applying these conditions during MS data analysis, TG2 was the 25th most abundant protein among the 413 RNA-binding proteins identified in the three samples (Supplementary Table 1) based on the number of detected unique peptides. In this list, only 20 proteins were newly identified RBPs according to the RBP2GO database, confirming the reliability of the applied experimental approach (Supplementary Table 2.). Notably, in the RBP2GO database, most of the RBPs were identified using tumour cell cultures with unlimited proliferation but less tissue specificity. Here, we used immortalised HUVEC cells and the newly recognised RBPs can be linked to endothelial functions such as angiogenesis, tube development, regulation of blood vessel endothelial cell migration, and tissue development based on the STRING analysis, thus lending support to the relevance of our results (Supplementary Table 3.).

### HUVEC-derived and a synthetic 43-mer RNA binds to TG2

Since RNA sequence recognised by TG2 was initially not known, we used total RNA isolated from HUVEC as a bait for a magnetic RNA-protein pull-down experiment after 3’ terminal labelling with desthiobiotin conjugated cytidine by the Pierce RNA 3’ End Desthiobiotinylation Kit containing T4 DNA ligase. This labelled total RNA pulled down recombinant human TG2 (Fig. 2A). The 43-mer RNA molecule, provided by the manufacturer for positive control of the labelling reaction as a good substrate for T4 DNA ligase, also interacted with the recombinant TG2 protein while non-specific binding of proteins to the magnetic beads was not detected in the absence of immobilised RNA. This 43-mer RNA has a strong secondary structure predicted by RNAstructure v6.4 (Fig. 2B).

**Figure 2.**
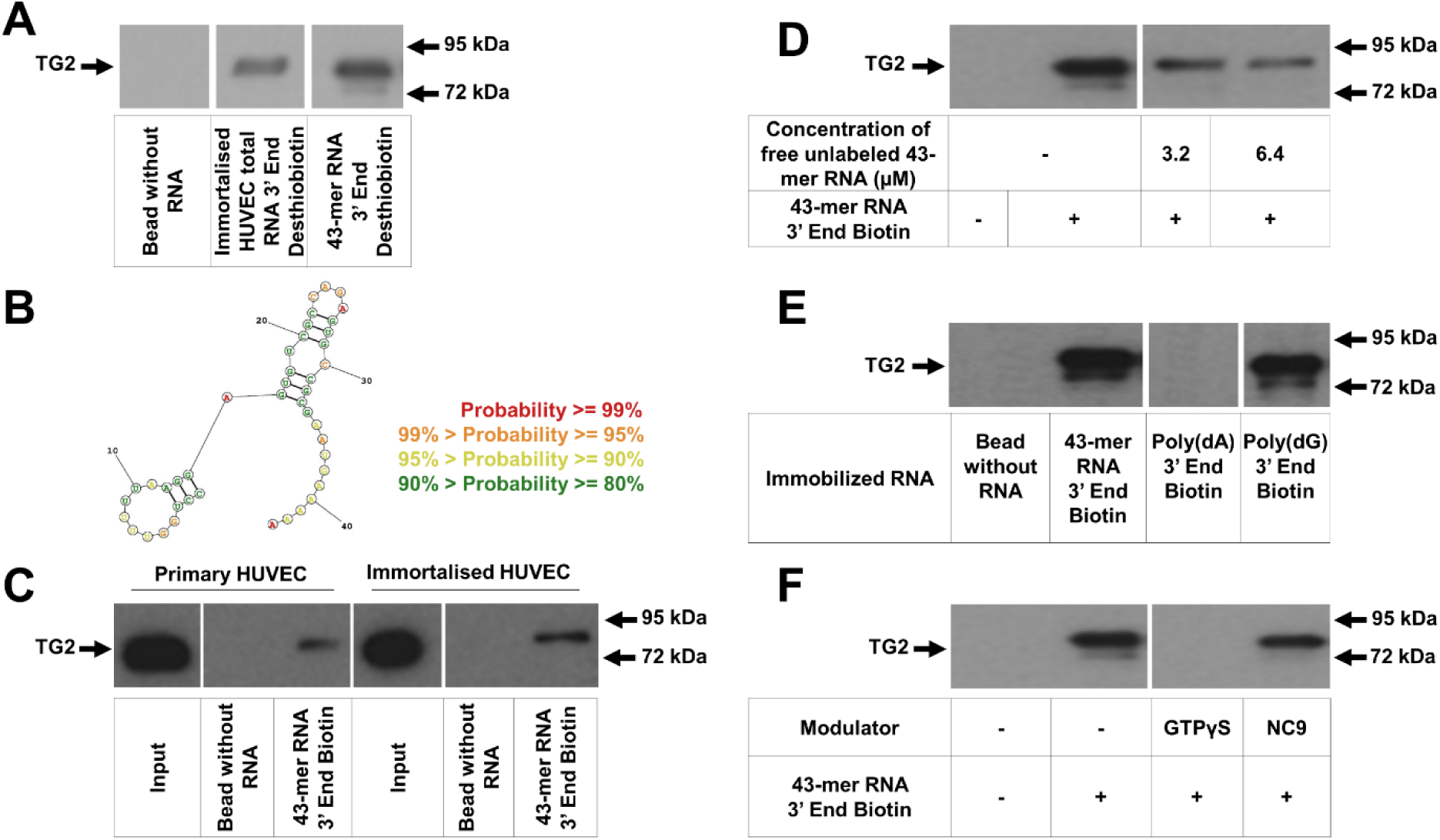
Magnetic RNA-protein pull-down experiments demonstrated the interaction between nucleic acids and TG2. (A) Both 3’ end desthiobiotinylated total RNA from HUVEC and a 43-mer RNA can bind recombinant human TG2. (B) The predicted secondary structure of the 43-mer RNA by RNAstructure v6.4 (Mathews lab). (C) 3’ end biotinylated 43-mer RNA can pull down endogenous TG2 from primary HUVEC and immortalised HUVEC protein extracts (each containing 200 µg protein). The input cell extracts contained 0.5 µg of protein. (D) Unlabelled 43-mer RNA competes with immobilised 3’ end biotinylated 43-mer RNA for TG2 binding. (E) Human recombinant TG2 can also bind to the 3’ end of the biotinylated homo-oligomeric poly(dG), but not to poly(dA). (F) In the presence of GTPγS, closed TG2 cannot, while NC9 stabilised open TG2 can bind to RNA.

In subsequent experiments, we used this 43-mer RNA in commercially available 3’ end biotin labelled form. First, it served as bait to pull down endogenous TG2 from the HUVEC protein extract. Fig. 2C demonstrates that endogenous TG2, present in either primary or immortalised HUVEC, can be co-precipitated with the applied 43-mer RNA molecule, providing further evidence for the existence of TG2-RNA complexes in HUVEC.

### Characterisation of RNA-binding property of recombinant TG2

In order to confirm the specificity of the interaction between TG2 and RNA, we performed a competition test in which recombinant TG2 was co-incubated with unlabelled 43-mer RNA and immobilised 43-mer RNA-coated magnetic beads (Fig. 2D). In the presence of increasing concentrations of unlabelled 43-mer RNA, the amount of pulled-down recombinant TG2 was decreased, suggesting that the ribonucleic acid is responsible for TG2 binding.

To determine the potential overlap between the RNA and GTP/GDP or ATP/ADP nucleotide binding sites of TG2 and to test the potential sequence and structural requirements of binding at the nucleic acid side, single-stranded poly(dG) and poly(dA) homo-oligomeric deoxy-nucleic acid were used in pull-down experiments. In the case of poly(dA), no interaction was detected, but poly(dG) could interact with TG2 (Fig. 2E). These findings suggest that the guanine base and its binding site in TG2 could be involved, at least partially, in the RNA-TG2 interaction. Thus, both the sequence and the secondary structure motifs of the nucleic acids seem to be important for binding to TG2, while negative charges are not sufficient for the interaction.

To investigate whether the conformation of TG2 influences its RNA binding, an RNA-protein magnetic pull-down experiment was performed in the presence of GTPγS, which is known to shift TG2 to a closed conformation or upon NC9 TG2 inhibitor treatment, which keeps the enzyme in an open structure (Akbar et al., 2017). GTPγS prevented the interaction of TG2 with the 43-mer RNA, while NC9 treatment did not influence RNA binding (Fig. 2F), suggesting that the RNA-binding surface on the TG2 protein is available only in its open form and covered in the closed, nucleotide-bound form.

### Determination of TG2 binding affinity for the 43-mer RNA molecule

Biolayer interferometry is a label-free optical technique for detecting and quantifying molecular interactions real-time based on the change in reflectance at the sensor surface upon immobilisation and complex formation. We used the Blitz instrument for our measurements, which can measure only one sample at a time and has a drop and a tube sample holder, resulting in a shift between the steps of measurement at the change of sample holders.

To determine real-time binding curves and affinity between TG2 and 43-mer RNA, increasing concentrations of recombinant tag-free TG2 were applied in the range of 2.5-200 µg/ml after similar loadings of the streptavidin biosensors with biotinylated 43-mer RNA (Fig. 3A). The raw binding curves were step-corrected and aligned for the association and dissociation phases (Fig. 3B). Increasing TG2 concentrations resulted in increasing saturation signals. The calculated K_D_ value was 88 nM, applying a 1:1 binding model.

**Figure 3.**
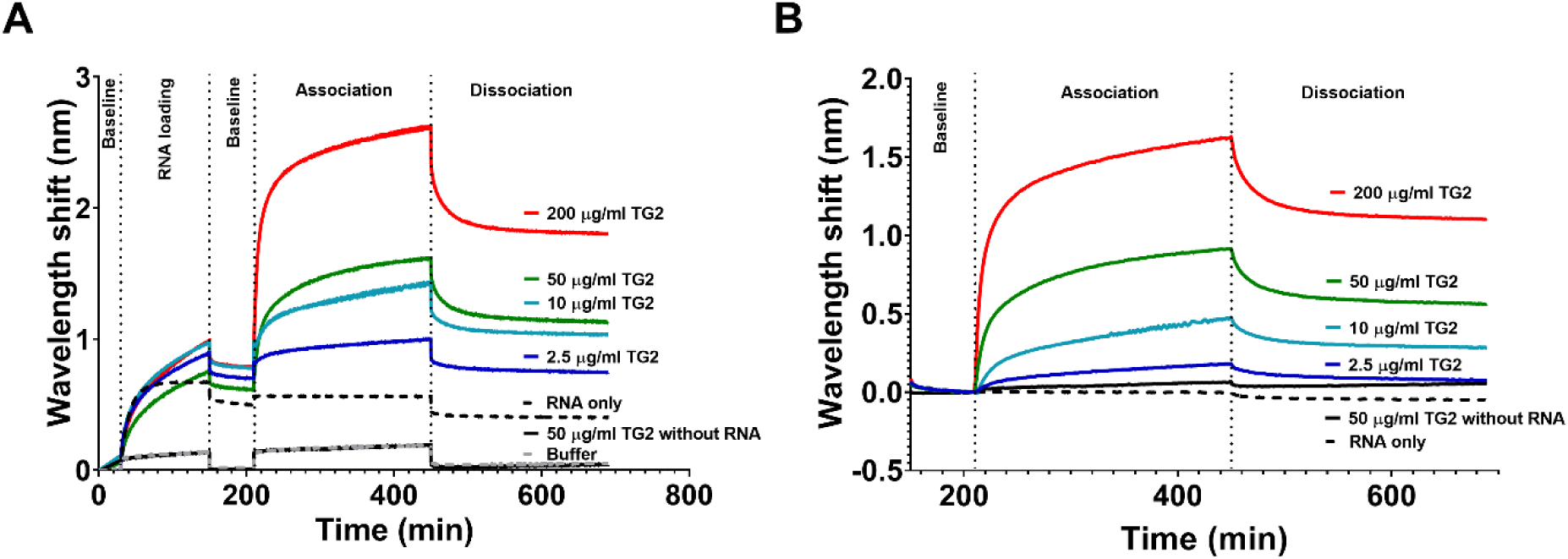
Real-time detection of TG2-RNA interactions using biolayer interferometry. (A) Raw binding curves of 43-mer RNA and recombinant expression tag-free human TG2. Representative curves based on at least two experiments generated by the Blitz instrument. (B) Step-corrected and aligned binding curves for the association and dissociation phases to determine the binding affinity between the 43-mer RNA and human TG2.

### Effect of 43-mer RNA on transglutaminase activities

Although RBPs can modulate RNA fate and function upon binding or releasing them, the interaction could also influence the biological activities of the RNA-binding protein. We examined the potential effect of 43-mer RNA binding to TG2 on two of its diverse transglutaminase activities. The RNA-binding affected neither transamidase nor isopeptidase activities, even at high RNA concentrations (Fig. 4).

**Figure 4.**
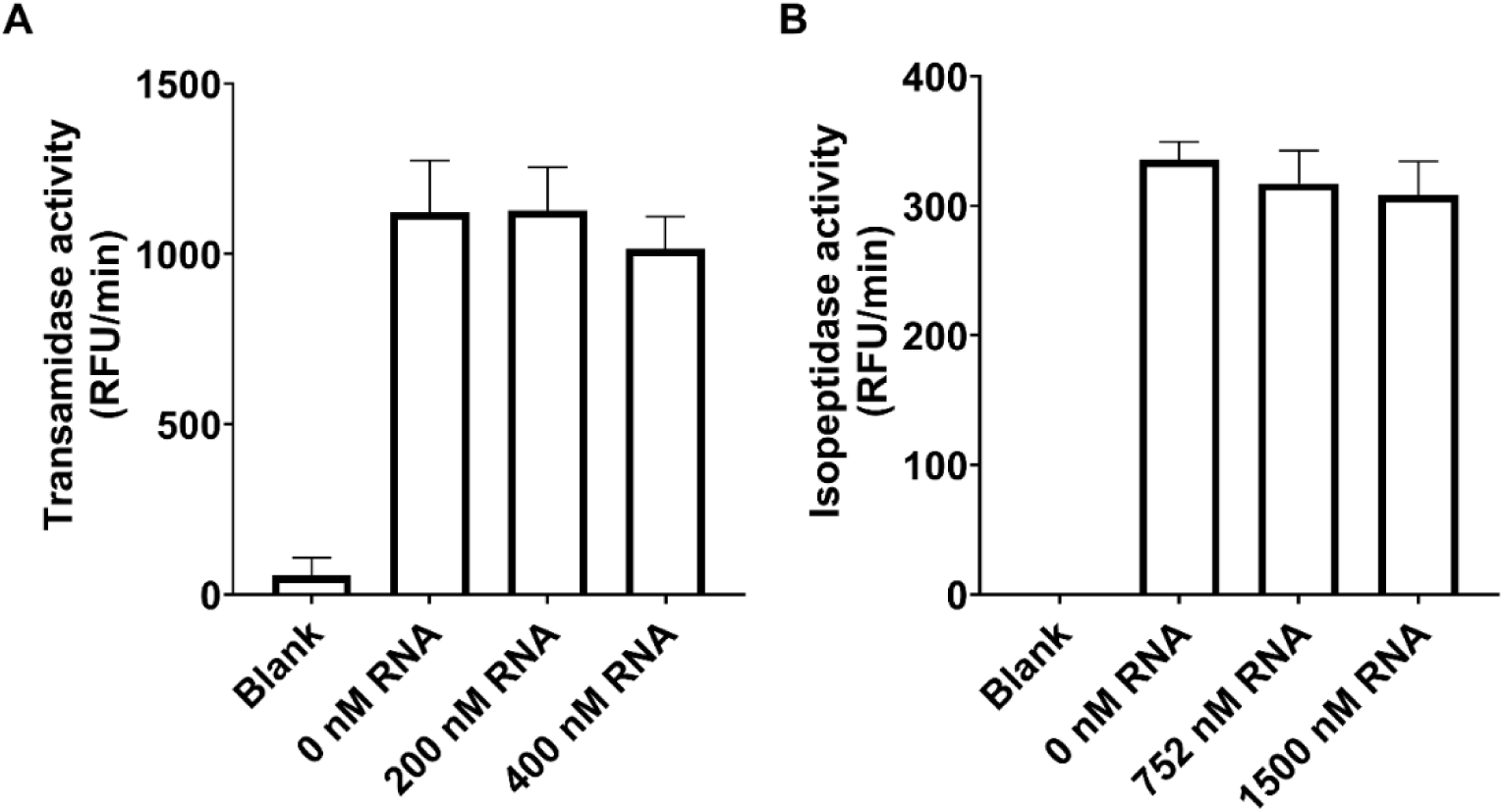
Effect of 43-mer RNA-binding on the transamidase (A) and isopeptidase (B) activity of TG2. The data are presented as the average ±SD of two independent experiments performed in triplicates. 200 nM or 150 nM recombinant (His)_6_-tagged TG2 was applied in the transamidase and isopeptidase assays, respectively. The blank contained 10 mM EDTA instead of calcium.

### *In silico* prediction of the RNA binding site in TG2

We performed *in silico* analyses to identify the RNA-binding site on the TG2 surface. The catRAPID algorithm (Agostini et al., 2013) calculates the RNA binding propensity of a protein; however, the human TG2 sequence was not predicted to bind RNA based on the calculated values. There is a broad spectrum of specific domains such as RNA-recognition domains (RRM), K-homology domains, and various zinc finger domains which, in combination with special, (*e.g.* RGG repeat) sequences, could be responsible for the high-affinity RNA binding of proteins (reviewed in Corley et al., 2020; Ottoz & Berchowitz 2020; Zeke et al., 2022). TG2 does not have these typical domains, as its nucleotide- and calcium-binding sites are also non-canonical (Király et al., 2011). Intrinsically disordered regions (IDRs) are frequently combined to RNA-binding motifs and are themselves involved in RNA-binding (Ottoz & Berchowitz, 2020; Zeke et al., 2022). TG2 has several predicted IDRs identified in an earlier publication (Thangaraju et al., 2017). Structured RNA-binding motifs frequently contain Arg, Lys, and His residues, and Arg and Ser repeats, aromatic amino acids, Tyr and Phe, and YGG repeats are also enriched residues in RNA-binding IDRs (Ottoz & Berchowitz, 2020). These can support direct interactions with RNA by Π-stacking, H-bonding and electrostatic interactions or just simply provide flexibility to the peptide chain. Merging the multifarious information using VMD visualisation software, the TG2 structure was analysed for potential RNA binding sites (Fig. 5). First, we selected stretches in which positively charged amino acids are frequent (similarly to known RNA binding motifs) and could overlap with predicted IDR regions. These surface patches were coloured on the visualised models of TG2 open and closed conformation (Fig. 5). As Poly(dG) can bind to TG2, and the known GTP-binding residues are on the surface in the open form but are partially hidden in the closed form, not binding RNA, residues in these surface patches could be involved in RNA binding. In Fig. 5, the blue-coloured residues (K173, F174, I175, K176, N177, I178, P179, N181, and Q186) are either responsible for the GTP binding (K173) or are in a proximal position potentially contributing to the nucleic acid-binding on the core domain. The C-terminal part of the GTP/GDP binding site and surrounding potential RNA-binding residues were coloured green (R580, Y583, R476, R478, Q481, S482, M483, H494, and V479). There are some IDRs with a high ratio of residues frequently involved in RNA binding coloured by orange (R433, D434, E435, R436, E437, H441, K444, P446, E447, G448, S449, and R453) and brown (K598, Q599, K600, R601, and K602). Finally, the purple colour labels basic residues in and around the nuclear export signal because this region has several typical RNA-binding residues (H658, H662, K663, N667, 672K, K674, and K677). Considering that the closed conformation abolished RNA binding, suggesting that the interacting surface is located somewhere on that surface of TG2, which is hidden in the closed conformation, brown and orange could be excluded because they are in surface positions in both conformations. After comparing the open and closed TG2 structures, we predicted that partially, the GTP-binding site on the core domain (Fig. 5, blue) and positively charged residues on the C-terminal barrel domain (Fig. 5, purple) could be responsible for the RNA binding, because these residues are not completely accessible in the closed conformation, suggesting that they are partially hidden in the closed TG2. Further studies using truncated and mutated TG2 variants or proteomics approaches are needed to determine the exact RNA-binding site of TG2.

**Figure 5.**
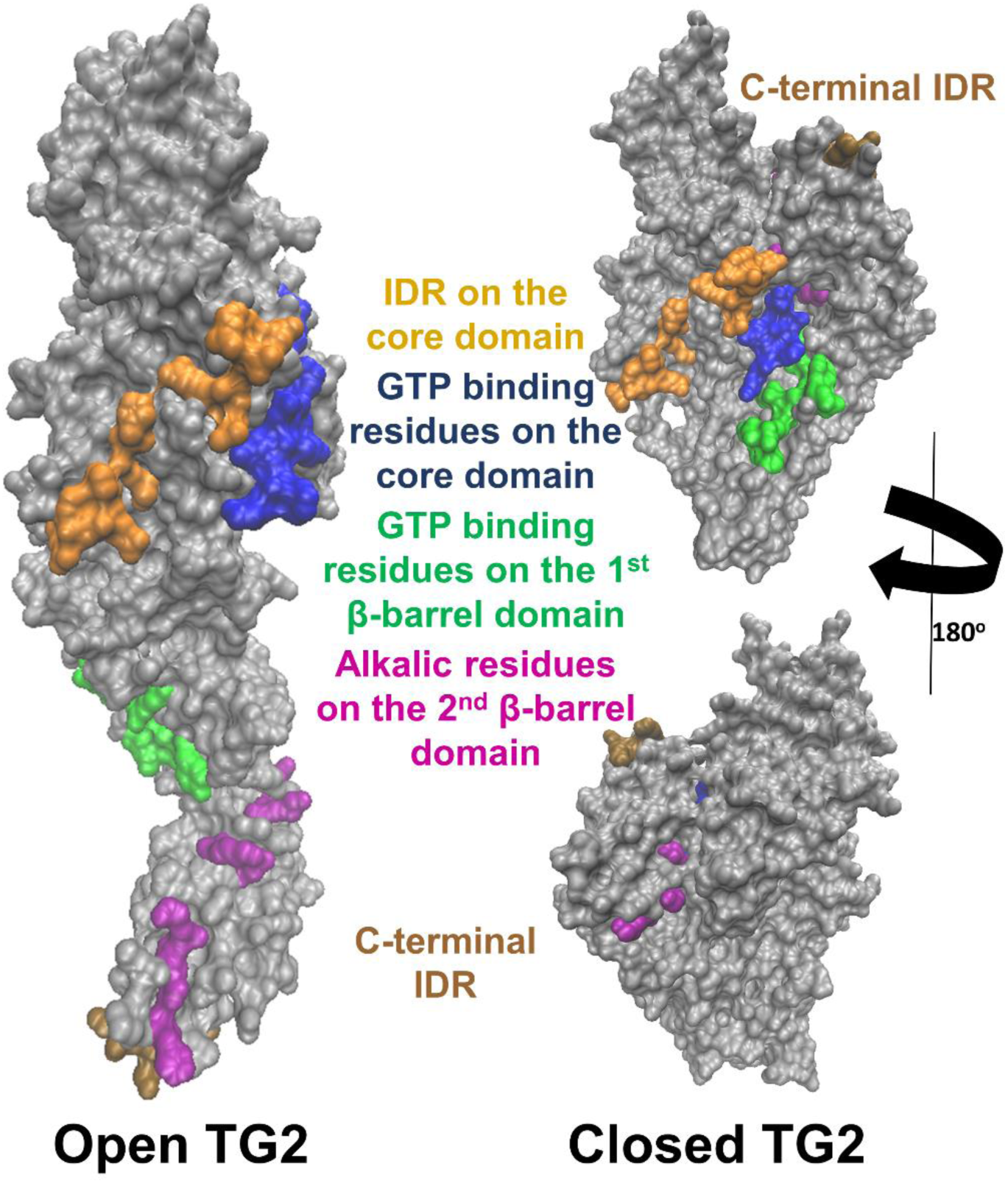
Structural representation of potential RNA-binding regions in the TG2 protein. The left image shows the open TG2 conformation (PDB 2q3z), while the right upper and lower images represent the closed TG2 conformation from the opposite side view (PDB 1kv3). Number of labelled amino acids: Green (corresponding to the C-terminal region of the GTP binding site): R580, Y583, R476, R478, Q481, S482, M483, H494, and V479. Blue (correlates with the core domain part of the GTP binding site): K173, F174, I175, K176, N177, I178, P179, N181, and Q186. Orange (extended IDR with alkalic residues on the core domain): R433, D434, E435, R436, E437, H441, K444, P446, E447, G448, S449, and R453. Purple (surface localised alkalic residues covered in the closed form): H658, H662, K663, N667, K672, K674, and K677. Brown (IDR close to the C-terminus): K598, Q599, K600, R601, and K602.

## Discussion

RBPs are heterogeneous proteins that interact with RNA molecules to regulate various aspects of gene expression and post-transcriptional processes, such as splicing, transport and modulation of mRNA translation. There are slightly different methods for identifying RBPs, but they always start with the formation of covalent bonds using UV light to stabilise the RNA-RBP complexes in cells. Then, these covalent complexes are isolated without enrichment using a phase separation protocol or with enrichment, for example, using oligo-dT resin to isolate mRNAs and their interactome. After RNA cleavage, mass spectrometry or specific antibodies are used to identify the released proteins (Castello et al., 2012). In recent years, the number of identified RBP candidates in the RBP2GO database reached 6100 (accessed on March 20, 2024). The available techniques also make the identification of bound RNA sequences possible (Caudron-Herger et al., 2021).

In the RBP2GO database, there are records about the presence of transglutaminases in RNA-protein complexes. In addition to human TG2, which was isolated from U2OS cells by OOPS as a potential RBP (Queiroz et al., 2019), human TG1 and TG3 were observed in 3 and 2 RBP-related studies, respectively, using HEK293, Jurkat, HuH-7 cell lines. Other human transglutaminases, such as TG4-7 and F13A, have not been identified as RNA-binding proteins. By performing an extensive search, another study was found that tried to identify RBPs recognising and binding to hairpins in pre/pri-miRNAs: here, a pull-down based experiment, when various miRNA hairpins were immobilised, led to the identification of TG2 as an RBP (Treiber et al., 2017). In contrast to the previous studies, in which tumour cell lines were usually used to obtain large sample volumes for proteomics analyses, we isolated RBPs from an immortal HUVEC line that maintained its endothelial-like characteristics, expressed endothelial marker genes and exhibited functional properties of HUVEC cells upon cytokine treatment (Shaw et al., 2021).

Our experimental results strongly support the existence of TG2-RNA interaction in HUVEC cells since we were able to demonstrate it by magnetic pull-down assay between the total RNA isolated from the HUVEC cells and bacterially expressed recombinant TG2, and vice versa when a 43-mer RNA was immobilised and it fished out endogenous TG2 from the HUVEC protein extract. This interaction likely requires a particular RNA sequence or secondary structure because, despite the ATP/ADP binding property of TG2, poly(dA) did not interact with TG2. To define the potential biological relevance of this interaction, we performed a BLAST search in the Ensemble database for the entire 43-mer RNA sequence, but this search did not yield a proper hit with reasonable similarity. When the 43-mer RNA secondary structure was predicted (Fig. 2B), we observed a loop similar to an AU-rich element (ARE) and another loop with bulges providing potential structural elements for interacting with an RBP. Further study is needed to determine the exact sequence and structure of TG2-interacting RNA molecules; this research is in progress in our laboratory.

Biolayer interferometry measurements were performed to determine the strength of the interaction between TG2 and the 43-mer RNA. Concentration-dependent binding curves were detected when biotinylated 43-mer RNA was immobilised on the streptavidin sensor surface. The determined K_D_ value was 88 nM. Protein-RNA binding affinities were compared by Nithin and coworkers (Nithin et al., 2019), who reported a relatively broad range of binding affinities. For example, between iron regulatory protein 1 and ferritin H IRE RNA, the K_D_ is 45 pM, while for nuclear RNA export factor 1 protein and CTE RNA the K_D_ is 90 nM, and several K_D_ values are in the micromolar range. We have determined the K_D_ between TG2 and a biologically irrelevant RNA molecule in the nanomolar range, suggesting that the natural cellular interacting RNA could have an even higher affinity towards TG2.

One potential biological relevance of RNA binding to a protein is riboregulation, when RNA binding controls the biological activities and functions of a protein. As an example, the reversible activity of cytoplasmic serine hydroxymethyltransferase 1 (SHMT1) is directed towards serine when the mRNA of SHMT2 competes with the polyglutamylated folate substrate (Guiducci et al., 2019). TG2 catalyses diverse activities, and the bound RNA could cause a switch between these. We have tested the effect of 43-mer RNA binding on the opposite transamidase and isopeptidase activities of TG2 and have not observed significant effects. Although the 43-mer RNA did not influence transglutaminase activities in the applied experimental conditions, the *in vivo* riboregulation of TG2 by cellular RNAs when it utilises natural cellular substrates cannot be ruled out. RNA binding may be responsible for the fine regulation of TG2, potentially influencing its Ca^2+^ sensitivity, GTP/GDP binding, and sensitivity against reducing/oxidising effects. RNA binding can affect the localisation of proteins, modulating their interactions with transport proteins (Hentze et al., 2018). The ratio of TG2 in the nucleus and cytoplasm depends on a non-canonical nuclear export signal (Kojima et al., 2012), which overlaps with the potential RNA binding surface described above, raising the possibility that RNAs can participate in the nuclear-cytoplasmic transport of TG2. Recognition and silencing of natural TG2-binding RNAs can help clarify these issues.

What is the biological relevance of the RNA binding capability of TG2? Since RBPs are involved in all steps of RNA regulation, and TG2 is present in the cytoplasm and nucleus, it could regulate posttranscriptional gene expression at any step, contributing to various processes. In mammalian mRNAs, AU-rich elements (ARE), which are partially similar to the 43-mer RNA, direct mRNA toward rapid degradation in unstimulated cells, ensuring that proteins transcribed from these mRNAs are expressed at very low levels (Lourou et al., 2019). mRNAs with poly-U regions encode proteins that are essential for inflammation and cell proliferation, and their expression levels are raised when necessary, usually for a short time. For example, in activated T-cells, the TIA-1 protein binds to the ARE of mRNA encoding the Fas receptor, promoting the synthesis of its pro-apoptotic form (Lourou et al., 2019). This finding also demonstrated that RBPs are vital for the appropriate progression of inflammation and apoptosis, preventing autoimmunity or tumour disease, in which the participation of TG2 has already been confirmed (Lourou et al., 2019; Eckert et al., 2014). Although, TG2 has several known activities to modulate transcription factors affecting the expression of specific genes (Tatsukawa et al., 2016), its newly confirmed RNA-binding ability suggests that it may act at various levels in these processes. For example, the binding of SHMT1 to the mRNA of SHMT2 silences the translation from that mRNA (Guiducci et al., 2019).

Previously, in a few cell lines, Treiber and coworkers attempted to catalogue the RBPs affecting the maturation and formation of miRNAs (Treiber et al., 2017). According to this study, TG2 can interact with 17 microRNAs, potentially influencing their production. However, Treiber’s study did not examine TG2-bound miRNAs in detail, but based on the GeneCards database, several miRNAs interacting with TG2 play a role in endothelial cell functions. For example, miR-21 negatively regulates endothelial cell proliferation, miR-29c, miR-106b, and miR-214 negatively regulate angiogenesis, while miR-31, miR-200a and miR-214 can act on blood vessel endothelial cell migration. These data raise the possibility that TG2 fundamentally affects endothelial cell functions as an RNA-binding protein regulating the maturation and functions of some critically important miRNAs.

Lastly, it should also be considered that TG2 could be externalised via extracellular vesicles (Zemskov et al., 2011). Like other extracellular and sorting proteins, TG2 may participate in externalising and transporting various RNA molecules (Shinde A et al., 2020). Vesicle-transported RNAs are involved in several physiological processes and can also contribute to the development of pathological conditions.

Taken together, our results highlighted a previously unknown feature of the TG2 protein, demonstrating that TG2 interacts with RNA molecules in HUVEC, and this interaction is affected by the sequence and secondary structure of the RNA. The RNA binding affinity of TG2 is high, and the interaction could be even more potent with natural ligands and in the cellular environment. There are several pieces of information suggesting the biological significance of the RNA-binding property of TG2, but further studies are needed to reveal the details of the related processes. Further characterisation of the RNA-binding property of transglutaminase 2 may reveal mechanisms in regulating its multi-functionality.

## Materials and Methods

All materials were purchased from Sigma-Aldrich (Munich, Germany) unless otherwise indicated.

### Cell extract preparation

Primary or immortalised HUVEC were cultured in T75 flasks until confluency, then trypsin-EDTA treatment for 5 minutes at 37°C was applied to detach the cells, which were washed 2 times with ice-cold sterile PBS. The cell pellet was resuspended in lysis buffer (25 mM Tris-HCl, 150 mM NaCl, 1 mM EDTA, 5% glycerol, 0.5% NP-40, pH 7.4, 1 mM PMSF, 1% protease inhibitor cocktail, 1% PhosSTOP (Roche, #04 906 845 001). Then, cell lysates were sonicated with 5 impulses using a merged sonicator tip, and cell debris was separated by spin centrifugation. The supernatant was used as cell extract for further experiments after determination of the protein concentration using Bio-Rad Protein Assay Dye Reagent Concentrate (Bio-Rad, Hercules, CA, USA).

### Isolation of RNA

For total RNA isolation, trypsinised and double-washed cells in PBS were resuspended in 1 ml TRI Reagent (Molecular Research Centre, Cincinnati OH, United States, cat#TR 118), and total RNA was isolated based on previously published protocol (Klusóczki et al., 2019).

### Orthogonal Organic Phase Separation

RNA-binding proteins were isolated using the orthogonal organic phase separation (OOPS) technique, according to the previous publication in Nature Protocols (Queiroz et al., 2019). Briefly, for each isolation, HUVEC were grown in two 10 cm Petri dishes. After two washes with PBS, the cells were irradiated with 400 mJ/cm^2^ UV light to fixate the RNA-protein complexes by covalent cross-linking in the cells (GS Gene Linker UV Chamber, Bio-Rad); except for the control samples which were not UV-treated. Then, the cells were lysed with TRI reagent. The homogenized lysates were transferred into Eppendorf tubes. For phase separation, chloroform was added (at a 1:5 ratio) and the aqueous and organic phases were carefully removed. The interphase (approximately 100 µl) was washed three times by repeating the addition of TRI reagent and chloroform. After washing, the interphases containing the UV cross-linked RNA-protein complexes were precipitated using methanol (9:1, vol/vol, methanol:sample). The pellet was resuspended in 0.1 M Tris-HCl, pH 8.5, containing 1% SDS.

After sonication (15 min, using 30 sec sonication/rest cycles in a cold water bath sonicator; BD ProbeTec ET, 2510E-DTH, Becton Dickinson, Maryland, USA), and consequent denaturation (at 95°C for 5 min), samples were treated with 5 µg RNase A (Thermo Scientific, Waltham, MA USA, #EN0531) at 37°C for 4 hours. To separate the released proteins from the complexes, again, TRI Reagent and chloroform were added, and equal volumes of the organic phases were applied for methanol precipitation from each sample. After washing (80% ethanol), the pellets were resolved in 100-100 µl of 0.1 M Tris-HCl buffer, pH 8.5, containing 1% SDS. For Western blotting, the same sample volumes were prepared by adding 6x denaturation buffer containing 50 mM dithiotreitol (DTT).

### LC-MS/MS analysis

After OOPS separation, protein samples were loaded on a 10% polyacrylamide gel and run until the beginning of the separating gel. After Coomassie blue staining (PageBlue Protein Staining Solution, Thermo Scientific, #24620), the visualised unseparated protein band was cut out from the gel and subjected to mass spectrometric analysis.

Protein identification based on the peptide sequences (MS/MS spectra) was performed by MaxQuant 1.6.2.10. search engine (Cox et al., 2008.) using the SwissProt/UniProt database. The results were imported into the Scaffold 5.0 program (Proteome Software, Inc., Portland, OR, USA). To prevent the loss of any potential RBPs, non-FDR filtering mode and one unique peptide per protein criterion were used, but the MS/MS spectrum of each peptide was verified by the presence of at least four consequent b or y fragment ions similar to our earlier publication (Csobán-Szabó et al., 2021).

### Production of recombinant TG2 proteins

The human recombinant N-terminal (His)_6_-tagged TG2 protein (containing valine at position 224; pET-30 Ek/LIC-TG2) was produced in *E. coli Rosetta2(DE3)* and purified by Ni-NTA affinity chromatography as described previously (Király et al., 2009). To determine the binding affinity between TG2 and 43-mer RNA a purified tag-free TG2 protein was also produced by cleaving the (His)_6_-tag by applying the TEV protease based on an earlier publication (Biri et al., 2016.) Protein concentrations were measured using Bio-Rad Protein Assay Dye Reagent Concentrate, and the purity was checked using SDS-PAGE followed by Coomassie blue staining or Western blotting with a monoclonal anti-TG2 antibody (CUB7402, Thermo, #MA5-12736).

### RNA-protein pull-down experiment

RNA-protein interactions were detected using Pierce RNA 3’ End Desthiobiotinylation Kit and Pierce Magnetic RNA-Protein Pull-Down Kit (Thermo Scientific, #20163), according to the manufacturer’s instructions.

493 µg total RNA, isolated from immortalised HUVEC cells and 50 pmol 43-mer RNA molecule included in the kit were desthiobiotinylated by ligating labelled cytidine to the nucleic acids. The ligation reaction mixture was incubated overnight at 16°C to enhance ligation effectivity. The precipitation, washing and redissolution steps of the labelled RNA molecules were performed according to the manufacturer’s instructions.

All of the labelled RNA samples were bound to 50 µl streptavidin magnetic bead suspension provided in the kit. For the development of RNA-protein complexes, after washing with 100 µl 20 mM Tris-HCl, pH 7.5, human (His)_6_-tagged recombinant TG2 was added to the RNA-loaded magnetic beads (using the suggested master mix) and it was incubated for 1 hour at room temperature. After elimination of the unbound protein, the beads were washed three times and the RNA-bound protein was eluted using 6x denaturation buffer and heat denaturation at 100°C for 5 minutes. The eluted samples were analysed using Western blotting after 10% SDS-PAGE. In the case of the first RNA-protein pull-down experiments, when desthiobiotinylated RNAs were used, 28 µg of human recombinant TG2 was applied for the binding. Then, 50 pmol of synthetic biotinylated RNA or single-stranded DNA (poly(dA) or poly(dG)) were loaded onto the Pierce Streptavidin Magnetic Bead (Thermo Scientific, #20164), and an experimentally normalised 5 µg of human recombinant TG2 was used (this resulted in readily detectable signal after the pull-down).

To check the conformation dependency of TG2’s RNA binding, the recombinant protein was preincubated for 15-20 minutes at room temperature with 0.1 mM NC9 modulator for the open conformation or TG2 protein and the RNA loaded beads were incubated in the presence of 0.5 mM GTPγS to form the closed conformation. The sources of the applied oligonucleotides are listed in Table 1.

**Table 1.**
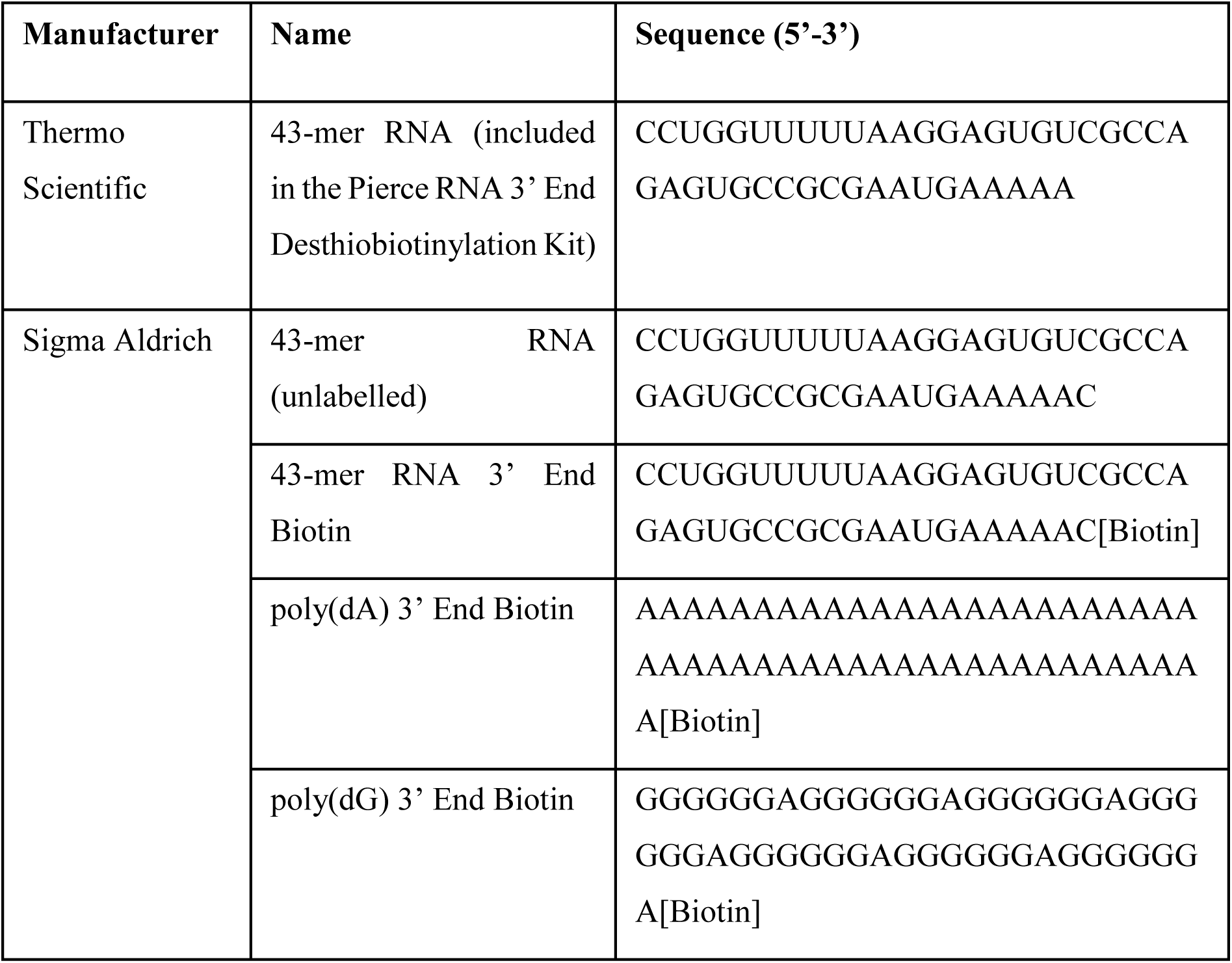
RNA and DNA oligonucleotides used in the study.

### Western blot

SDS-PAGE (10%) was followed by a semidry blotting method using PVDF membrane (Millipore Immobilon-P membrane, 0.45 µm). For blocking 5 (*m/v*) % low-fat milk powder containing TTBS was used for 1 hour at room temperature or overnight. Primary monoclonal transglutaminase-2 antibody (CUB7402, Thermo Fisher #MA5-12736, 1/15000) was diluted in 0.5 (*m/v*) % low-fat milk powder in TTBS and incubated for 1 hour at room temperature or overnight at 4°C. After four washes (15 minutes with TTBS), the secondary goat-anti-mouse IgG/HRP antibody (Advansta #R-05071-500, 1/10000) was diluted in 0.5 (*m/v*) % low-fat milk powder in TTBS and incubated for one hour at room temperature. After the washing steps, the Western blot was visualised using WesternBrite ECL HRP Substrate (Advansta).

### Kinetic transamidase assay

Transamidase activity, which involves the incorporation of dansyl-cadaverine into N, N’-dimethylated casein, was measured based on a previously published procedure (Király et al., 2016). The reaction mixture contained 50 mM Tris-HCl, pH 8.0, 0.2 mM dansyl-cadaverine, 2 mg/ml N, N’-dimethylated casein, 3 mM DTT, and 10 mM CaCl_2_ or 10 mM EDTA. The blank contained 10 mM EDTA instead of CaCl_2_. The assay was started by adding TG2 (200 nM final concentration in a 100 µl reaction volume). The activity was calculated based on the linear part of the increasing fluorescence (Ex/Em: 360/530 nm, at 37°C).

### Fluorescent isopeptidase assay

Isopeptidase activity was determined using the A102 fluorescence substrate (Zedira, Darmstadt, Germany), according to an earlier publication (Király et al., 2016). Briefly, transglutaminases cleave the isopeptide bound in the synthesised substrate, releasing the dark quencher (2,4-dinitrophenyl). The final concentrations in the reaction mixture were 50 mM MOPS buffer, pH 6.8, 10 mM CaCl_2_, 100 mM NaCl, 0.1% (w/v) PEG8000, 50 μM A102 and 3 mM DTT. The blank contained 10 mM EDTA instead of CaCl_2_. The reaction was started by adding TG2 (150 nM final concentration in a 200 µl volume) and was monitored at 37°C by a BIOTEK Synergy 4 or H1 multiple reader (Ex/Em: 318/413 nm).

### Biolayer interferometry

Biolayer interferometry is an optical technique for measuring molecular interactions in real-time and determining the kinetics and affinity of these interactions. The binding kinetics were determined using the BLItz System (PALL FortéBio, Fremont, CA, USA). Streptavidin biosensors (PALL FortéBio) were loaded with 10 µg/ml of biotinylated 43-mer RNA in binding buffer (0.02 M Tris-HCl, pH 7.5 containing 0.05 M NaCl, 2 mM MgCl_2_ and 0.1% Tween 20) using the drop sample holder. The purified (His)_6_-tag-free recombinant human TG2 was also applied using the drop sample holder, and the measurements were made in duplicates. For other steps tube sample holder was used, which generated a shift in the curve between the experimental steps. We applied a step correction to eliminate these shifts between subsequent experimental steps. The dissociation constant (K_D_) was determined by BLItz Pro 1.2.1.5. software using the association and dissociation aligned curves applying a “Global”, 1:1 kinetic model. For control experiments signal of RNA loading, without adding any protein, and then only protein association in the absence of immobilised RNA were recorded using separate sensor tips.

### Bioinformatics analyses

The presence of identified RNA-binding proteins from HUVEC was checked in the RBP2GO database (rbp2go.dkfz.de; Caudron-Herger et al., 2021). The 20 newly revealed potential RNA-binding proteins were analysed using the STRING database (string-db.org). The most enriched GO biological functions are listed based on the FDR (false discovery rate) values. The secondary structure of the 43-mer RNA was predicted using RNAstructure v6.4 (Mathews lab, rna.urmc.rochester.edu/RNAstructureWeb). The potential RNA-binding ability of TG2 was estimated by the catRAPID online tool (Armaos et al., 2021). The visual representation of TG2 was made using VMD 1.9.4a53 (www.ks.uiuc.edu/Research/vmd/; Humphrey et al., 1996).

## Author Contributions

Conceptualisation, R.K.; methodology, B.C., I.R.K-S. and R.K.; software, A.R.C. and B.C; formal analysis, visualisation, B.C., A.R.C. and R.K.; investigation, B.C., A.R.C., I.R.K-S. and R.K.; data curation, B.C.; writing—original draft preparation, R.K. and B.C..; writing—review and editing, R.K., I.R.K-S. and L.F.; funding acquisition, resources, R.K. and L.F. All authors have read and agreed to the published version of the manuscript.

## Acknowledgements

We thank János Mótyán for the critical reading and suggestions for the manuscript preparation and Jennifer Nagy for technical assistance in the laboratory (Department of Biochemistry and Molecular Biology, University of Debrecen). We appreciate the helpful comments and service of Gergő Kalló and the Proteomics Core Facility (mass spectrometry analysis), respectively, as well as Szilárd Póliska and the Genomic Medicine and Bioinformatics Core Facility (RNAseq analysis), in the Department of Biochemistry and Molecular Biology, University of Debrecen. This work was supported by the following grant: Hungarian Scientific Research Fund [NKFI OTKA K 129139].

## Data availability statement

Proteomics data will be available upon request from the corresponding author and after a one-year embargo in the ProteomeXchange database.

## Abbreviations

ARE: adenine-uridine rich element
Blitz: biolayer interferometry
EV: extracellular vesicle
HUVEC: human umbilical cord vein endothelial cell
IDR: intrinsically disordered region
OOPS: orthogonal organic phase separation
PDI: protein disulfide isomerase
RBP: RNA-binding protein
SHMT1: serine hydroxymethyltransferase
TG2: transglutaminase 2

