## Supplementary materials for "Transglutaminase 2 is an RNA-binding protein: Experimental verification and characterisation of a novel transglutaminase feature"

### Supplementary Figure

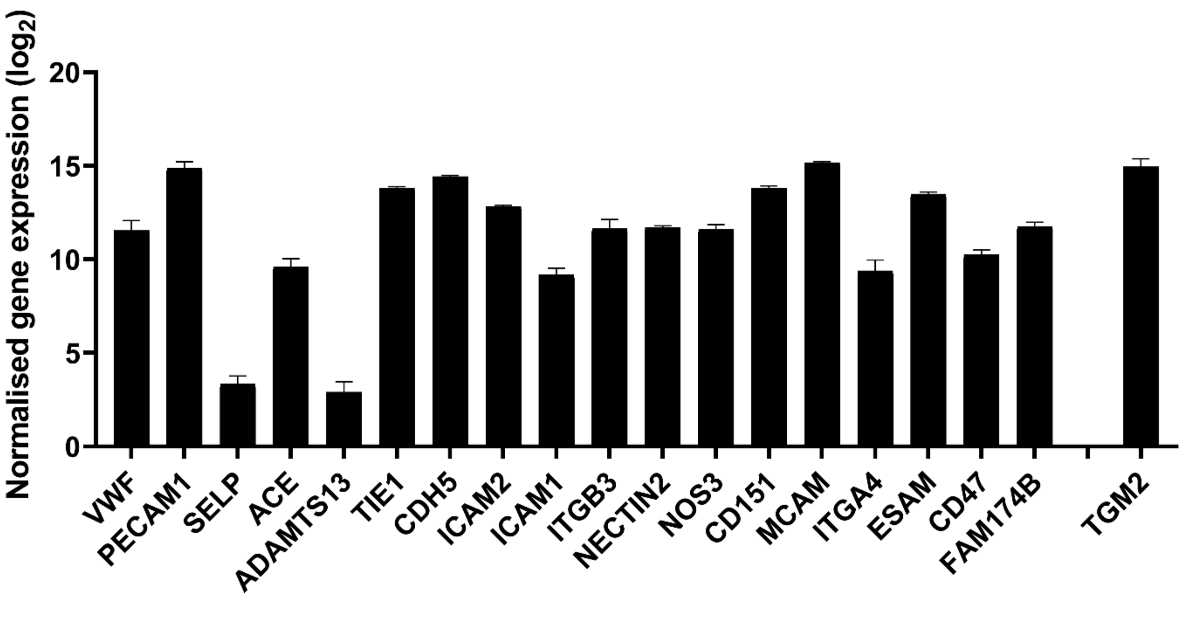


**Supplementary figure 1. Normalised expression of endothelial marker and TG2 genes in immortalised HUVEC cell line determined by mRNA sequencing**. Data are presented as average normalised gene expression ($log_2$) of three independent samples with SD.

**Supplementary table 1. Identified RNA-binding proteins in the immortalised HUVEC (n=3).**

| Identified Proteins (439) | Accession Number | Alternate ID | Molecular Weight | Peptidcount 1. sample | Peptidcount 2. sample | Peptidcount 3. sample | Peptidcount SUM |
|---|---|---|---|---|---|---|---|
| Myosin-9 OS=Homo sapiens OX=9606 GN=MYH9 PE=1 SV=4 | P35579 | MYH9 | 227 kDa | 44 | 69 | 61 | 174 |
| Plectin OS=Homo sapiens OX=9606 GN=PLEC PE=1 SV=3 | Q15149 | PLEC | 532 kDa | 30 | 54 | 56 | 140 |
| Vimentin OS=Homo sapiens OX=9606 GN=VIM PE=1 SV=4 | P08670 | VIM | 54 kDa | 39 | 39 | 36 | 114 |
| Filamin-A OS=Homo sapiens OX=9606 GN=FLNA PE=1 SV=4 | P21333 (+1) | FLNA | 281 kDa | 29 | 36 | 38 | 103 |
| Filamin-B OS=Homo sapiens OX=9606 GN=FLNB PE=1 SV=2 | O75369 (+2) | FLNB | 278 kDa | 25 | 39 | 36 | 100 |
| Annexin A2 OS=Homo sapiens OX=9606 GN=ANXA2 PE=1 SV=2 | P07355 | ANXA2 | 39 kDa | 23 | 25 | 35 | 83 |
| Neuroblast differentiation-associated protein AHNAK OS=Homo sapiens OX=9606 GN=AHNAK PE=1 SV=2 | Q09666 | AHNAK | 629 kDa | 15 | 35 | 25 | 75 |
| Prelamin-A/C OS=Homo sapiens OX=9606 GN=LMNA PE=1 SV=1 | P02545 | LMNA | 74 kDa | 23 | 28 | 22 | 73 |
| Talin-1 OS=Homo sapiens OX=9606 GN=TLN1 PE=1 SV=3 | Q9Y490 | TLN1 | 270 kDa | 15 | 25 | 25 | 65 |
| Actin, cytoplasmic 2 OS=Homo sapiens OX=9606 GN=ACTG1 PE=1 SV=1 | P63261 | ACTG1 | 42 kDa | 19 | 21 | 24 | 64 |
| Alpha-actinin-1 OS=Homo sapiens OX=9606 GN=ACTN1 PE=1 SV=2 | P12814 | ACTN1 | 103 kDa | 10 | 22 | 23 | 55 |
| Tubulin beta chain OS=Homo sapiens OX=9606 GN=TUBB PE=1 SV=2 | P07437 | TUBB | 50 kDa | 12 | 19 | 21 | 52 |
| Alpha-enolase OS=Homo sapiens OX=9606 GN=ENO1 PE=1 SV=2 | P06733 | ENO1 | 47 kDa | 17 | 13 | 18 | 48 |
| Cytoplasmic dynein 1 heavy chain 1 OS=Homo sapiens OX=9606 GN=DYNC1H1 PE=1 SV=5 | Q14204 | DYNC1H1 | 532 kDa | 9 | 23 | 15 | 47 |
| Pyruvate kinase PKM OS=Homo sapiens OX=9606 GN=PKM PE=1 SV=4 | P14618 | PKM | 58 kDa | 14 | 13 | 18 | 45 |
| Endoplasmic reticulum chaperone BiP OS=Homo sapiens OX=9606 GN=HSPA5 PE=1 SV=2 | P11021 | HSPA5 | 72 kDa | 13 | 15 | 16 | 44 |
| Filamin-C OS=Homo sapiens OX=9606 GN=FLNC PE=1 SV=3 | Q14315 (+1) | FLNC | 291 kDa | 11 | 16 | 16 | 43 |

| | | | | | | | |
|---|---|---|---|---|---|---|---|
| Elongation factor 2 OS=Homo sapiens OX=9606 GN=EEF2 PE=1 SV=4 | P13639 | EEF2 | 95 kDa | 12 | 13 | 17 | 42 |
| Clathrin heavy chain 1 OS=Homo sapiens OX=9606 GN=CLTC PE=1 SV=5 | Q00610 (+1) | CLTC | 192 kDa | 8 | 14 | 18 | 40 |
| Isoform 2 of Tubulin alpha-1A chain OS=Homo sapiens OX=9606 GN=TUBA1A | Q71U36-2 | TUBA1A | 46 kDa | 10 | 15 | 15 | 40 |
| Heat shock protein HSP 90-beta OS=Homo sapiens OX=9606 GN=HSP90AB1 PE=1 SV=4 | P08238 | HSP90AB1 | 83 kDa | 8 | 14 | 18 | 40 |
| Heat shock cognate 71 kDa protein OS=Homo sapiens OX=9606 GN=HSPA8 PE=1 SV=1 | P11142 | HSPA8 | 71 kDa | 9 | 13 | 15 | 37 |
| Glyceraldehyde-3-phosphate dehydrogenase OS=Homo sapiens OX=9606 GN=GAPDH PE=1 SV=3 | P04406 | GAPDH | 36 kDa | 13 | 12 | 12 | 37 |
| Annexin A1 OS=Homo sapiens OX=9606 GN=ANXA1 PE=1 SV=2 | P04083 | ANXA1 | 39 kDa | 10 | 10 | 15 | 35 |
| **Protein-glutamine gamma-glutamyltransferase 2 OS=Homo sapiens OX=9606 GN=TGM2 PE=1 SV=2** | **P21980** | **TGM2** | **77 kDa** | **8** | **12** | **13** | **33** |
| Transketolase OS=Homo sapiens OX=9606 GN=TKT PE=1 SV=3 | P29401 (+1) | TKT | 68 kDa | 10 | 10 | 11 | 31 |
| Heterogeneous nuclear ribonucleoprotein U OS=Homo sapiens OX=9606 GN=HNRNPU PE=1 SV=6 | Q00839 (+1) | HNRNPU | 91 kDa | 8 | 12 | 10 | 30 |
| Transitional endoplasmic reticulum ATPase OS=Homo sapiens OX=9606 GN=VCP PE=1 SV=4 | P55072 | VCP | 89 kDa | 9 | 10 | 10 | 29 |
| Isoform 2 of Ubiquitin-like modifier-activating enzyme 1 OS=Homo sapiens OX=9606 GN=UBA1 | P22314-2 | UBA1 | 114 kDa | 7 | 8 | 14 | 29 |
| Spectrin beta chain, non-erythrocytic 1 OS=Homo sapiens OX=9606 GN=SPTBN1 PE=1 SV=2 | Q01082 | SPTBN1 | 275 kDa | 7 | 11 | 10 | 28 |
| Alpha-actinin-4 OS=Homo sapiens OX=9606 GN=ACTN4 PE=1 SV=2 | O43707 | ACTN4 | 105 kDa | 5 | 12 | 10 | 27 |
| Protein disulfide-isomerase A3 OS=Homo sapiens OX=9606 GN=PDIA3 PE=1 SV=4 | P30101 | PDIA3 | 57 kDa | 11 | 7 | 9 | 27 |
| Elongation factor 1-alpha 1 OS=Homo sapiens OX=9606 GN=EEF1A1 PE=1 SV=1 | P68104 (+1) | EEF1A1 | 50 kDa | 9 | 9 | 8 | 26 |
| Fibronectin OS=Homo sapiens OX=9606 GN=FN1 PE=1 SV=5 | P02751 (+6) | FN1 | 272 kDa | 7 | 6 | 12 | 25 |

| | | | | | | | |
|---|---|---|---|---|---|---|---|
| Heterogeneous nuclear ribonucleoproteins A2/B1 OS=Homo sapiens OX=9606 GN=HNRNPA2B1 PE=1 SV=2 | P22626 | HNRNPA2B1 | 37 kDa | 6 | 10 | 9 | 25 |
| Myoferlin OS=Homo sapiens OX=9606 GN=MYOF PE=1 SV=1 | Q9NZM1 (+2) | MYOF | 235 kDa | 6 | 6 | 13 | 25 |
| Polyubiquitin-B OS=Homo sapiens OX=9606 GN=UBB PE=1 SV=1 | P0CG47 (+3) | UBB | 26 kDa | 9 | 9 | 7 | 25 |
| Basement membrane-specific heparan sulfate proteoglycan core protein OS=Homo sapiens OX=9606 GN=HSPG2 PE=1 SV=4 | P98160 | HSPG2 | 469 kDa | 6 | 6 | 11 | 23 |
| Nucleolin OS=Homo sapiens OX=9606 GN=NCL PE=1 SV=3 | P19338 | NCL | 77 kDa | 5 | 7 | 10 | 22 |
| Fructose-bisphosphate aldolase A OS=Homo sapiens OX=9606 GN=ALDOA PE=1 SV=2 | P04075 | ALDOA | 39 kDa | 6 | 9 | 6 | 21 |
| Moesin OS=Homo sapiens OX=9606 GN=MSN PE=1 SV=3 | P26038 | MSN | 68 kDa | 6 | 7 | 8 | 21 |
| Spectrin alpha chain, non-erythrocytic 1 OS=Homo sapiens OX=9606 GN=SPTAN1 PE=1 SV=3 | Q13813 (+2) | SPTAN1 | 285 kDa | 5 | 11 | 5 | 21 |
| Cofilin-1 OS=Homo sapiens OX=9606 GN=CFL1 PE=1 SV=3 | P23528 | CFL1 | 19 kDa | 5 | 8 | 8 | 21 |
| Unconventional myosin-Ic OS=Homo sapiens OX=9606 GN=MYO1C PE=1 SV=4 | O00159 | MYO1C | 122 kDa | 6 | 5 | 9 | 20 |
| Vinculin OS=Homo sapiens OX=9606 GN=VCL PE=1 SV=4 | P18206 (+1) | VCL | 124 kDa | 7 | 5 | 7 | 19 |
| Peroxiredoxin-1 OS=Homo sapiens OX=9606 GN=PRDX1 PE=1 SV=1 | Q06830 | PRDX1 | 22 kDa | 6 | 7 | 6 | 19 |
| Heterogeneous nuclear ribonucleoprotein R OS=Homo sapiens OX=9606 GN=HNRNPR PE=1 SV=1 | O43390 (+2) | HNRNPR | 71 kDa | 5 | 6 | 8 | 19 |
| Major vault protein OS=Homo sapiens OX=9606 GN=MVP PE=1 SV=4 | Q14764 | MVP | 99 kDa | 5 | 6 | 7 | 18 |
| Annexin A6 OS=Homo sapiens OX=9606 GN=ANXA6 PE=1 SV=3 | P08133 | ANXA6 | 76 kDa | 5 | 6 | 7 | 18 |
| Staphylococcal nuclease domain-containing protein 1 OS=Homo sapiens OX=9606 GN=SND1 PE=1 SV=1 | Q7KZF4 | SND1 | 102 kDa | 4 | 8 | 6 | 18 |
| Thrombospondin-1 OS=Homo sapiens OX=9606 GN=THBS1 PE=1 SV=2 | P07996 | THBS1 | 129 kDa | 6 | 5 | 6 | 17 |
| Cytoskeleton-associated protein 4 OS=Homo sapiens OX=9606 GN=CKAP4 PE=1 SV=2 | Q07065 | CKAP4 | 66 kDa | 4 | 7 | 6 | 17 |

| | | | | | | | |
|---|---|---|---|---|---|---|---|
| Heterogeneous nuclear ribonucleoprotein H OS=Homo sapiens OX=9606 GN=HNRNPH1 PE=1 SV=4 | P31943 | HNRNPH1 | 49 kDa | 5 | 5 | 7 | 17 |
| Microtubule-associated protein 4 OS=Homo sapiens OX=9606 GN=MAP4 PE=1 SV=3 | P27816 (+1) | MAP4 | 121 kDa | 7 | 7 | 2 | 16 |
| DNA-dependent protein kinase catalytic subunit OS=Homo sapiens OX=9606 GN=PRKDC PE=1 SV=3 | P78527 | PRKDC | 469 kDa | 4 | 5 | 7 | 16 |
| ATP-dependent 6-phosphofructokinase, platelet type OS=Homo sapiens OX=9606 GN=PFKP PE=1 SV=2 | Q01813 | PFKP | 86 kDa | 4 | 5 | 7 | 16 |
| Thioredoxin domain-containing protein 5 OS=Homo sapiens OX=9606 GN=TXNDC5 PE=1 SV=2 | Q8NBS9 | TXNDC5 | 48 kDa | 4 | 4 | 7 | 15 |
| Probable ATP-dependent RNA helicase DDX17 OS=Homo sapiens OX=9606 GN=DDX17 PE=1 SV=2 | Q92841 (+1) | DDX17 | 80 kDa | 3 | 6 | 6 | 15 |
| Chloride intracellular channel protein 1 OS=Homo sapiens OX=9606 GN=CLIC1 PE=1 SV=4 | O00299 | CLIC1 | 27 kDa | 6 | 4 | 5 | 15 |
| Stress-70 protein, mitochondrial OS=Homo sapiens OX=9606 GN=HSPA9 PE=1 SV=2 | P38646 | HSPA9 | 74 kDa | 7 | 4 | 4 | 15 |
| Annexin A5 OS=Homo sapiens OX=9606 GN=ANXA5 PE=1 SV=2 | P08758 | ANXA5 | 36 kDa | 5 | 5 | 5 | 15 |
| Lamin-B1 OS=Homo sapiens OX=9606 GN=LMNB1 PE=1 SV=2 | P20700 | LMNB1 | 66 kDa | 3 | 5 | 6 | 14 |
| Endoplasmin OS=Homo sapiens OX=9606 GN=HSP90B1 PE=1 SV=1 | P14625 | HSP90B1 | 92 kDa | 4 | 2 | 8 | 14 |
| WD repeat-containing protein 1 OS=Homo sapiens OX=9606 GN=WDR1 PE=1 SV=4 | O75083 | WDR1 | 66 kDa | 5 | 4 | 5 | 14 |
| Heterogeneous nuclear ribonucleoprotein K OS=Homo sapiens OX=9606 GN=HNRNPK PE=1 SV=1 | P61978 | HNRNPK | 51 kDa | 3 | 4 | 7 | 14 |
| Far upstream element-binding protein 2 OS=Homo sapiens OX=9606 GN=KHSRP PE=1 SV=4 | Q92945 | KHSRP | 73 kDa | 6 | 5 | 3 | 14 |
| Peptidyl-prolyl cis-trans isomerase A OS=Homo sapiens OX=9606 GN=PPIA PE=1 SV=2 | P62937 | PPIA | 18 kDa | 4 | 4 | 6 | 14 |
| Protein disulfide-isomerase A6 OS=Homo sapiens OX=9606 GN=PDIA6 PE=1 SV=1 | Q15084 (+4) | PDIA6 | 48 kDa | 7 | 3 | 4 | 14 |

| | | | | | | | |
|---|---|---|---|---|---|---|---|
| ATP-dependent RNA helicase A OS=Homo sapiens OX=9606 GN=DHX9 PE=1 SV=4 | Q08211 | DHX9 | 141 kDa | 3 | 5 | 5 | 13 |
| ATP synthase subunit alpha, mitochondrial OS=Homo sapiens OX=9606 GN=ATP5F1A PE=1 SV=1 | P25705 (+1) | ATP5F1A | 60 kDa | 3 | 5 | 5 | 13 |
| 60S ribosomal protein L4 OS=Homo sapiens OX=9606 GN=RPL4 PE=1 SV=5 | P36578 | RPL4 | 48 kDa | 3 | 4 | 6 | 13 |
| T-complex protein 1 subunit beta OS=Homo sapiens OX=9606 GN=CCT2 PE=1 SV=4 | P78371 | CCT2 | 57 kDa | 2 | 4 | 7 | 13 |
| Src substrate cortactin OS=Homo sapiens OX=9606 GN=CTTN PE=1 SV=2 | Q14247 (+1) | CTTN | 62 kDa | 4 | 5 | 4 | 13 |
| X-ray repair cross-complementing protein 6 OS=Homo sapiens OX=9606 GN=XRCC6 PE=1 SV=2 | P12956 | XRCC6 | 70 kDa | 6 | 3 | 4 | 13 |
| T-complex protein 1 subunit delta OS=Homo sapiens OX=9606 GN=CCT4 PE=1 SV=4 | P50991 (+1) | CCT4 | 58 kDa | 3 | 6 | 4 | 13 |
| Kinesin-1 heavy chain OS=Homo sapiens OX=9606 GN=KIF5B PE=1 SV=1 | P33176 | KIF5B | 110 kDa | 3 | 5 | 5 | 13 |
| Ras GTPase-activating-like protein IQGAP1 OS=Homo sapiens OX=9606 GN=IQGAP1 PE=1 SV=1 | P46940 | IQGAP1 | 189 kDa | 2 | 6 | 5 | 13 |
| Guanine nucleotide-binding protein G(i) subunit alpha-2 OS=Homo sapiens OX=9606 GN=GNAI2 PE=1 SV=3 | P04899 (+1) | GNAI2 | 40 kDa | 2 | 4 | 6 | 12 |
| BTB/POZ domain-containing protein KCTD12 OS=Homo sapiens OX=9606 GN=KCTD12 PE=1 SV=1 | Q96CX2 | KCTD12 | 36 kDa | 4 | 4 | 4 | 12 |
| Vigilin OS=Homo sapiens OX=9606 GN=HDLBP PE=1 SV=2 | Q00341 | HDLBP | 141 kDa | 3 | 6 | 3 | 12 |
| Calpain-2 catalytic subunit OS=Homo sapiens OX=9606 GN=CAPN2 PE=1 SV=6 | P17655 | CAPN2 | 80 kDa | 2 | 5 | 5 | 12 |
| Tubulin beta-6 chain OS=Homo sapiens OX=9606 GN=TUBB6 PE=1 SV=1 | Q9BUF5 | TUBB6 | 50 kDa | 1 | 4 | 7 | 12 |
| Polypyrimidine tract-binding protein 1 OS=Homo sapiens OX=9606 GN=PTBP1 PE=1 SV=1 | P26599 (+2) | PTBP1 | 57 kDa | 3 | 4 | 5 | 12 |
| Glucose-6-phosphate 1-dehydrogenase OS=Homo sapiens OX=9606 GN=G6PD PE=1 SV=4 | P11413 (+1) | G6PD | 59 kDa | 4 | 4 | 4 | 12 |
| L-lactate dehydrogenase A chain OS=Homo sapiens OX=9606 GN=LDHA PE=1 SV=2 | P00338 | LDHA | 37 kDa | 4 | 4 | 3 | 11 |

| | | | | | | | |
|---|---|---|---|---|---|---|---|
| Coronin-1B OS=Homo sapiens OX=9606 GN=CORO1B PE=1 SV=1 | Q9BR76 | CORO1B | 54 kDa | 3 | 4 | 4 | 11 |
| Neutral alpha-glucosidase AB OS=Homo sapiens OX=9606 GN=GANAB PE=1 SV=3 | Q14697 (+1) | GANAB | 107 kDa | 3 | 4 | 4 | 11 |
| Glutathione S-transferase P OS=Homo sapiens OX=9606 GN=GSTP1 PE=1 SV=2 | P09211 | GSTP1 | 23 kDa | 4 | 3 | 4 | 11 |
| T-complex protein 1 subunit theta OS=Homo sapiens OX=9606 GN=CCT8 PE=1 SV=4 | P50990 | CCT8 | 60 kDa | 4 | 2 | 5 | 11 |
| 40S ribosomal protein SA OS=Homo sapiens OX=9606 GN=RPSA PE=1 SV=4 | P08865 | RPSA | 33 kDa | 4 | 4 | 3 | 11 |
| Bifunctional glutamate/proline--tRNA ligase OS=Homo sapiens OX=9606 GN=EPRS1 PE=1 SV=5 | P07814 | EPRS1 | 171 kDa | 3 | 4 | 4 | 11 |
| Dolichyl-diphosphooligosaccharide--protein glycosyltransferase subunit 1 OS=Homo sapiens OX=9606 GN=RPN1 PE=1 SV=1 | P04843 | RPN1 | 69 kDa | 3 | 3 | 5 | 11 |
| ELAV-like protein 1 OS=Homo sapiens OX=9606 GN=ELAVL1 PE=1 SV=2 | Q15717 (+1) | ELAVL1 | 36 kDa | 2 | 3 | 6 | 11 |
| Keratin, type II cytoskeletal 1 OS=Homo sapiens OX=9606 GN=KRT1 PE=1 SV=6 | P04264 (+1) | KRT1 | 66 kDa | 4 | 3 | 3 | 10 |
| Cell surface glycoprotein MUC18 OS=Homo sapiens OX=9606 GN=MCAM PE=1 SV=2 | P43121 | MCAM | 72 kDa | 3 | 3 | 4 | 10 |
| Phosphoglycerate kinase 1 OS=Homo sapiens OX=9606 GN=PGK1 PE=1 SV=3 | P00558 | PGK1 | 45 kDa | 2 | 3 | 5 | 10 |
| Fatty acid synthase OS=Homo sapiens OX=9606 GN=FASN PE=1 SV=3 | P49327 | FASN | 273 kDa | 2 | 2 | 6 | 10 |
| Nestin OS=Homo sapiens OX=9606 GN=NES PE=1 SV=2 | P48681 | NES | 177 kDa | 2 | 4 | 4 | 10 |
| Myosin light polypeptide 6 OS=Homo sapiens OX=9606 GN=MYL6 PE=1 SV=2 | P60660 (+1) | MYL6 | 17 kDa | 2 | 3 | 5 | 10 |
| Synaptic vesicle membrane protein VAT-1 homolog OS=Homo sapiens OX=9606 GN=VAT1 PE=1 SV=2 | Q99536 | VAT1 | 42 kDa | 3 | 2 | 5 | 10 |
| Keratin, type II cytoskeletal 7 OS=Homo sapiens OX=9606 GN=KRT7 PE=1 SV=5 | P08729 (+1) | KRT7 | 51 kDa | 3 | 4 | 3 | 10 |

| | | | | | | | |
|---|---|---|---|---|---|---|---|
| Heterogeneous nuclear ribonucleoprotein F OS=Homo sapiens OX=9606 GN=HNRNPF PE=1 SV=3 | P52597 | HNRNPF | 46 kDa | 2 | 2 | 6 | 10 |
| Multifunctional protein ADE2 OS=Homo sapiens OX=9606 GN=PAICS PE=1 SV=3 | P22234 (+1) | PAICS | 47 kDa | 3 | 4 | 3 | 10 |
| Ras-interacting protein 1 OS=Homo sapiens OX=9606 GN=RASIP1 PE=1 SV=1 | Q5U651 | RASIP1 | 103 kDa | 1 | 2 | 7 | 10 |
| D-3-phosphoglycerate dehydrogenase OS=Homo sapiens OX=9606 GN=PHGDH PE=1 SV=4 | O43175 | PHGDH | 57 kDa | 3 | 3 | 4 | 10 |
| Isoform 2 of Protein flightless-1 homolog OS=Homo sapiens OX=9606 GN=FLII | Q13045-2 | FLII | 138 kDa | 2 | 4 | 4 | 10 |
| Ribosome-binding protein 1 OS=Homo sapiens OX=9606 GN=RRBP1 PE=1 SV=5 | Q9P2E9 | RRBP1 | 152 kDa | 5 | 2 | 2 | 9 |
| 60 kDa heat shock protein, mitochondrial OS=Homo sapiens OX=9606 GN=HSPD1 PE=1 SV=2 | P10809 | HSPD1 | 61 kDa | 2 | 3 | 4 | 9 |
| Protein disulfide-isomerase OS=Homo sapiens OX=9606 GN=P4HB PE=1 SV=3 | P07237 | P4HB | 57 kDa | 2 | 3 | 4 | 9 |
| EH domain-containing protein 2 OS=Homo sapiens OX=9606 GN=EHD2 PE=1 SV=2 | Q9NZN4 | EHD2 | 61 kDa | 2 | 3 | 4 | 9 |
| Plastin-3 OS=Homo sapiens OX=9606 GN=PLS3 PE=1 SV=4 | P13797 (+2) | PLS3 | 71 kDa | 2 | 3 | 4 | 9 |
| Coatomer subunit alpha OS=Homo sapiens OX=9606 GN=COPA PE=1 SV=2 | P53621 (+1) | COPA | 138 kDa | 2 | 3 | 4 | 9 |
| Transcription intermediary factor 1-beta OS=Homo sapiens OX=9606 GN=TRIM28 PE=1 SV=5 | Q13263 | TRIM28 | 89 kDa | 3 | 4 | 2 | 9 |
| Importin subunit beta-1 OS=Homo sapiens OX=9606 GN=KPNB1 PE=1 SV=2 | Q14974 | KPNB1 | 97 kDa | 1 | 4 | 4 | 9 |
| Isoform 2 of Tropomyosin alpha-3 chain OS=Homo sapiens OX=9606 GN=TPM3 | P06753-2 (+2) | TPM3 | 29 kDa | 2 | 3 | 4 | 9 |
| Transaldolase OS=Homo sapiens OX=9606 GN=TALDO1 PE=1 SV=2 | P37837 | TALDO1 | 38 kDa | 4 | 1 | 4 | 9 |
| Matrin-3 OS=Homo sapiens OX=9606 GN=MATR3 PE=1 SV=2 | P43243 | MATR3 | 95 kDa | 2 | 2 | 5 | 9 |
| C-1-tetrahydrofolate synthase, cytoplasmic OS=Homo sapiens OX=9606 GN=MTHFD1 PE=1 SV=4 | P11586 | MTHFD1 | 102 kDa | 4 | 4 | 1 | 9 |

| | | | | | | | |
|---|---|---|---|---|---|---|---|
| Heat shock protein beta-1 OS=Homo sapiens OX=9606 GN=HSPB1 PE=1 SV=2 | P04792 | HSPB1 | 23 kDa | 3 | 2 | 4 | 9 |
| Microtubule-associated protein 1B OS=Homo sapiens OX=9606 GN=MAP1B PE=1 SV=2 | P46821 | MAP1B | 271 kDa | 2 | 3 | 4 | 9 |
| 60S ribosomal protein L6 OS=Homo sapiens OX=9606 GN=RPL6 PE=1 SV=3 | Q02878 | RPL6 | 33 kDa | 2 | 3 | 4 | 9 |
| Isoform 3 of Zinc finger protein 185 OS=Homo sapiens OX=9606 GN=ZNF185 | O15231-3 | ZNF185 | 74 kDa | 2 | 5 | 2 | 9 |
| F-actin-capping protein subunit alpha-2 OS=Homo sapiens OX=9606 GN=CAPZA2 PE=1 SV=3 | P47755 (+1) | CAPZA2 | 33 kDa | 3 | 2 | 4 | 9 |
| Transgelin-2 OS=Homo sapiens OX=9606 GN=TAGLN2 PE=1 SV=3 | P37802 | TAGLN2 | 22 kDa | 0 | 4 | 5 | 9 |
| Heterogeneous nuclear ribonucleoprotein M OS=Homo sapiens OX=9606 GN=HNRNPM PE=1 SV=3 | P52272 (+1) | HNRNPM | 78 kDa | 2 | 3 | 3 | 8 |
| Heterogeneous nuclear ribonucleoproteins C1/C2 OS=Homo sapiens OX=9606 GN=HNRNPC PE=1 SV=4 | P07910 (+1) | HNRNPC | 34 kDa | 1 | 3 | 4 | 8 |
| Catenin delta-1 OS=Homo sapiens OX=9606 GN=CTNND1 PE=1 SV=1 | O60716 (+7) | CTNND1 | 108 kDa | 2 | 3 | 3 | 8 |
| Splicing factor, proline- and glutamine-rich OS=Homo sapiens OX=9606 GN=SFPQ PE=1 SV=2 | P23246 (+1) | SFPQ | 76 kDa | 2 | 3 | 3 | 8 |
| 60S ribosomal protein L8 OS=Homo sapiens OX=9606 GN=RPL8 PE=1 SV=2 | P62917 | RPL8 | 28 kDa | 1 | 4 | 3 | 8 |
| PDZ and LIM domain protein 1 OS=Homo sapiens OX=9606 GN=PDLIM1 PE=1 SV=4 | O00151 | PDLIM1 | 36 kDa | 2 | 2 | 4 | 8 |
| Coatomer subunit gamma-1 OS=Homo sapiens OX=9606 GN=COPG1 PE=1 SV=1 | Q9Y678 | COPG1 | 98 kDa | 2 | 4 | 2 | 8 |
| Heat shock protein HSP 90-alpha OS=Homo sapiens OX=9606 GN=HSP90AA1 PE=1 SV=5 | P07900 (+1) | HSP90AA1 | 85 kDa | 2 | 2 | 4 | 8 |
| ATP-dependent RNA helicase DDX3X OS=Homo sapiens OX=9606 GN=DDX3X PE=1 SV=3 | O00571 | DDX3X | 73 kDa | 1 | 3 | 4 | 8 |
| Rab GDP dissociation inhibitor beta OS=Homo sapiens OX=9606 GN=GDI2 PE=1 SV=2 | P50395 | GDI2 | 51 kDa | 2 | 0 | 6 | 8 |

| | | | | | | | |
|---|---|---|---|---|---|---|---|
| CAD protein OS=Homo sapiens OX=9606 GN=CAD PE=1 SV=3 | P27708 | CAD | 243 kDa | 2 | 1 | 5 | 8 |
| Tubulin beta-4B chain OS=Homo sapiens OX=9606 GN=TUBB4B PE=1 SV=1 | P68371 | TUBB4B | 50 kDa | 2 | 3 | 3 | 8 |
| Protein AHNAK2 OS=Homo sapiens OX=9606 GN=AHNAK2 PE=1 SV=2 | Q8IVF2 (+1) | AHNAK2 | 617 kDa | 0 | 4 | 4 | 8 |
| Isoform 6 of Unconventional myosin-VI OS=Homo sapiens OX=9606 GN=MYO6 | Q9UM54-6 | MYO6 | 149 kDa | 2 | 3 | 3 | 8 |
| Cytoplasmic FMR1-interacting protein 1 OS=Homo sapiens OX=9606 GN=CYFIP1 PE=1 SV=1 | Q7L576 | CYFIP1 | 145 kDa | 1 | 3 | 4 | 8 |
| Heterogeneous nuclear ribonucleoprotein A1 OS=Homo sapiens OX=9606 GN=HNRNPA1 PE=1 SV=5 | P09651 (+2) | HNRNPA1 | 39 kDa | 3 | 3 | 2 | 8 |
| Cysteine--tRNA ligase, cytoplasmic OS=Homo sapiens OX=9606 GN=CARS1 PE=1 SV=3 | P49589 (+1) | CARS1 | 85 kDa | 3 | 2 | 3 | 8 |
| Inositol 1,4,5-trisphosphate receptor type 3 OS=Homo sapiens OX=9606 GN=ITPR3 PE=1 SV=2 | Q14573 | ITPR3 | 304 kDa | 2 | 3 | 3 | 8 |
| Sodium/potassium-transporting ATPase subunit alpha-1 OS=Homo sapiens OX=9606 GN=ATP1A1 PE=1 SV=1 | P05023 (+2) | ATP1A1 | 113 kDa | 2 | 3 | 3 | 8 |
| Nuclear mitotic apparatus protein 1 OS=Homo sapiens OX=9606 GN=NUMA1 PE=1 SV=2 | Q14980 | NUMA1 | 238 kDa | 1 | 3 | 4 | 8 |
| EH domain-containing protein 4 OS=Homo sapiens OX=9606 GN=EHD4 PE=1 SV=1 | Q9H223 | EHD4 | 61 kDa | 3 | 2 | 3 | 8 |
| Eukaryotic translation initiation factor 3 subunit C-like protein OS=Homo sapiens OX=9606 GN=EIF3CL PE=1 SV=1 | B5ME19 (+2) | EIF3CL | 105 kDa | 3 | 2 | 3 | 8 |
| Valine--tRNA ligase OS=Homo sapiens OX=9606 GN=VARS1 PE=1 SV=4 | P26640 | VARS1 | 140 kDa | 0 | 3 | 5 | 8 |
| Isoform 2 of Glycine--tRNA ligase OS=Homo sapiens OX=9606 GN=GARS1 | P41250-2 | GARS1 | 78 kDa | 3 | 3 | 2 | 8 |
| T-complex protein 1 subunit gamma OS=Homo sapiens OX=9606 GN=CCT3 PE=1 SV=4 | P49368 | CCT3 | 61 kDa | 2 | 2 | 3 | 7 |
| Programmed cell death 6-interacting protein OS=Homo sapiens OX=9606 GN=PDCD6IP PE=1 SV=1 | Q8WUM4 | PDCD6IP | 96 kDa | 1 | 3 | 3 | 7 |

| | | | | | | | |
|---|---|---|---|---|---|---|---|
| ATP-citrate synthase OS=Homo sapiens OX=9606 GN=ACLY PE=1 SV=3 | P53396 (+1) | ACLY | 121 kDa | 1 | 3 | 3 | 7 |
| Integrin beta-1 OS=Homo sapiens OX=9606 GN=ITGB1 PE=1 SV=2 | P05556 | ITGB1 | 88 kDa | 1 | 2 | 4 | 7 |
| Coronin-1C OS=Homo sapiens OX=9606 GN=CORO1C PE=1 SV=1 | Q9ULV4 (+2) | CORO1C | 53 kDa | 3 | 2 | 2 | 7 |
| Eukaryotic initiation factor 4A-I OS=Homo sapiens OX=9606 GN=EIF4A1 PE=1 SV=1 | P60842 | EIF4A1 | 46 kDa | 2 | 1 | 4 | 7 |
| Phosphoglycerate mutase 1 OS=Homo sapiens OX=9606 GN=PGAM1 PE=1 SV=2 | P18669 | PGAM1 | 29 kDa | 1 | 3 | 3 | 7 |
| Galectin-1 OS=Homo sapiens OX=9606 GN=LGALS1 PE=1 SV=2 | P09382 | LGALS1 | 15 kDa | 2 | 2 | 3 | 7 |
| LIM and SH3 domain protein 1 OS=Homo sapiens OX=9606 GN=LASP1 PE=1 SV=2 | Q14847 | LASP1 | 30 kDa | 2 | 3 | 2 | 7 |
| Fermitin family homolog 3 OS=Homo sapiens OX=9606 GN=FERMT3 PE=1 SV=1 | Q86UX7 (+1) | FERMT3 | 76 kDa | 1 | 1 | 5 | 7 |
| Profilin-1 OS=Homo sapiens OX=9606 GN=PFN1 PE=1 SV=2 | P07737 | PFN1 | 15 kDa | 2 | 2 | 3 | 7 |
| EF-hand domain-containing protein D2 OS=Homo sapiens OX=9606 GN=EFHD2 PE=1 SV=1 | Q96C19 | EFHD2 | 27 kDa | 3 | 3 | 1 | 7 |
| T-complex protein 1 subunit zeta OS=Homo sapiens OX=9606 GN=CCT6A PE=1 SV=3 | P40227 | CCT6A | 58 kDa | 2 | 1 | 4 | 7 |
| Serine/threonine-protein kinase N1 OS=Homo sapiens OX=9606 GN=PKN1 PE=1 SV=2 | Q16512 (+1) | PKN1 | 104 kDa | 0 | 4 | 3 | 7 |
| Ubiquitin carboxyl-terminal hydrolase 5 OS=Homo sapiens OX=9606 GN=USP5 PE=1 SV=2 | P45974 (+1) | USP5 | 96 kDa | 2 | 3 | 2 | 7 |
| 14-3-3 protein zeta/delta OS=Homo sapiens OX=9606 GN=YWHAZ PE=1 SV=1 | P63104 | YWHAZ | 28 kDa | 3 | 1 | 3 | 7 |
| 40S ribosomal protein S8 OS=Homo sapiens OX=9606 GN=RPS8 PE=1 SV=2 | P62241 | RPS8 | 24 kDa | 0 | 3 | 4 | 7 |
| SH3 and multiple ankyrin repeat domains protein 3 OS=Homo sapiens OX=9606 GN=SHANK3 PE=1 SV=3 | Q9BYB0 | SHANK3 | 185 kDa | 2 | 3 | 2 | 7 |
| EH domain-containing protein 1 OS=Homo sapiens OX=9606 GN=EHD1 PE=1 SV=2 | Q9H4M9 | EHD1 | 61 kDa | 2 | 2 | 3 | 7 |

| | | | | | | | |
|---|---|---|---|---|---|---|---|
| Arf-GAP with Rho-GAP domain, ANK repeat and PH domain-containing protein 1 OS=Homo sapiens OX=9606 GN=ARAP1 PE=1 SV=3 | Q96P48 (+1) | ARAP1 | 162 kDa | 3 | 2 | 2 | 7 |
| 60S ribosomal protein L5 OS=Homo sapiens OX=9606 GN=RPL5 PE=1 SV=3 | P46777 | RPL5 | 34 kDa | 1 | 3 | 2 | 6 |
| Zyxin OS=Homo sapiens OX=9606 GN=ZYX PE=1 SV=1 | Q15942 | ZYX | 61 kDa | 0 | 3 | 3 | 6 |
| X-ray repair cross-complementing protein 5 OS=Homo sapiens OX=9606 GN=XRCC5 PE=1 SV=3 | P13010 | XRCC5 | 83 kDa | 1 | 2 | 3 | 6 |
| Dihydropyrimidinase-related protein 2 OS=Homo sapiens OX=9606 GN=DPYSL2 PE=1 SV=1 | Q16555 | DPYSL2 | 62 kDa | 2 | 2 | 2 | 6 |
| Probable ATP-dependent RNA helicase DDX5 OS=Homo sapiens OX=9606 GN=DDX5 PE=1 SV=1 | P17844 (+1) | DDX5 | 69 kDa | 2 | 2 | 2 | 6 |
| V-type proton ATPase catalytic subunit A OS=Homo sapiens OX=9606 GN=ATP6V1A PE=1 SV=2 | P38606 | ATP6V1A | 68 kDa | 1 | 2 | 3 | 6 |
| T-complex protein 1 subunit alpha OS=Homo sapiens OX=9606 GN=TCP1 PE=1 SV=1 | P17987 | TCP1 | 60 kDa | 3 | 2 | 1 | 6 |
| Heterogeneous nuclear ribonucleoprotein L OS=Homo sapiens OX=9606 GN=HNRNPL PE=1 SV=2 | P14866 | HNRNPL | 64 kDa | 1 | 2 | 3 | 6 |
| Constitutive coactivator of PPAR-gamma-like protein 1 OS=Homo sapiens OX=9606 GN=FAM120A PE=1 SV=2 | Q9NZB2 (+2) | FAM120A | 122 kDa | 1 | 2 | 3 | 6 |
| Elongation factor 1-gamma OS=Homo sapiens OX=9606 GN=EEF1G PE=1 SV=3 | P26641 | EEF1G | 50 kDa | 1 | 2 | 3 | 6 |
| T-complex protein 1 subunit epsilon OS=Homo sapiens OX=9606 GN=CCT5 PE=1 SV=1 | P48643 | CCT5 | 60 kDa | 2 | 1 | 3 | 6 |
| Galactokinase OS=Homo sapiens OX=9606 GN=GALK1 PE=1 SV=1 | P51570 | GALK1 | 42 kDa | 2 | 2 | 2 | 6 |
| Copine-1 OS=Homo sapiens OX=9606 GN=CPNE1 PE=1 SV=1 | Q99829 | CPNE1 | 59 kDa | 2 | 1 | 3 | 6 |
| Histidine--tRNA ligase, cytoplasmic OS=Homo sapiens OX=9606 GN=HARS1 PE=1 SV=2 | P12081 | HARS1 | 57 kDa | 3 | 1 | 2 | 6 |
| Actin, cytoplasmic 1 OS=Homo sapiens OX=9606 GN=ACTB PE=1 SV=1 | P60709 | ACTB | 42 kDa | 2 | 2 | 2 | 6 |
| Tryptophan--tRNA ligase, cytoplasmic OS=Homo sapiens OX=9606 GN=WARS1 PE=1 SV=2 | P23381 | WARS1 | 53 kDa | 2 | 0 | 4 | 6 |

| | | | | | | | |
|---|---|---|---|---|---|---|---|
| Non-POU domain-containing octamer-binding protein OS=Homo sapiens OX=9606 GN=NONO PE=1 SV=4 | Q15233 | NONO | 54 kDa | 3 | 2 | 1 | 6 |
| L-lactate dehydrogenase B chain OS=Homo sapiens OX=9606 GN=LDHB PE=1 SV=2 | P07195 | LDHB | 37 kDa | 2 | 1 | 3 | 6 |
| Isoform 5 of Septin-9 OS=Homo sapiens OX=9606 GN=SEPTIN9 | Q9UHD8-5 | SEPTIN9 | 65 kDa | 2 | 1 | 3 | 6 |
| RNA-binding motif protein, X chromosome OS=Homo sapiens OX=9606 GN=RBMX PE=1 SV=3 | P38159 | RBMX | 42 kDa | 3 | 2 | 1 | 6 |
| Drebrin-like protein OS=Homo sapiens OX=9606 GN=DBNL PE=1 SV=1 | Q9UJU6 | DBNL | 48 kDa | 2 | 3 | 1 | 6 |
| 60S ribosomal protein L24 OS=Homo sapiens OX=9606 GN=RPL24 PE=1 SV=1 | P83731 | RPL24 | 18 kDa | 1 | 2 | 3 | 6 |
| Sequestosome-1 OS=Homo sapiens OX=9606 GN=SQSTM1 PE=1 SV=1 | Q13501 | SQSTM1 | 48 kDa | 2 | 2 | 2 | 6 |
| Fascin OS=Homo sapiens OX=9606 GN=FSCN1 PE=1 SV=3 | Q16658 | FSCN1 | 55 kDa | 2 | 2 | 2 | 6 |
| 60S ribosomal protein L13 OS=Homo sapiens OX=9606 GN=RPL13 PE=1 SV=4 | P26373 | RPL13 | 24 kDa | 2 | 2 | 2 | 6 |
| GTPase-activating protein and VPS9 domain-containing protein 1 OS=Homo sapiens OX=9606 GN=GAPVD1 PE=1 SV=2 | Q14C86 (+5) | GAPVD1 | 165 kDa | 2 | 2 | 2 | 6 |
| 14-3-3 protein theta OS=Homo sapiens OX=9606 GN=YWHAQ PE=1 SV=1 | P27348 | YWHAQ | 28 kDa | 1 | 2 | 3 | 6 |
| Leucine-rich repeat-containing protein 47 OS=Homo sapiens OX=9606 GN=LRRC47 PE=1 SV=1 | Q8N1G4 | LRRC47 | 63 kDa | 1 | 2 | 3 | 6 |
| Nucleoplasmin-3 OS=Homo sapiens OX=9606 GN=NPM3 PE=1 SV=3 | O75607 | NPM3 | 19 kDa | 2 | 2 | 2 | 6 |
| Aminopeptidase N OS=Homo sapiens OX=9606 GN=ANPEP PE=1 SV=4 | P15144 | ANPEP | 110 kDa | 0 | 2 | 3 | 5 |
| Ribonuclease inhibitor OS=Homo sapiens OX=9606 GN=RNH1 PE=1 SV=2 | P13489 | RNH1 | 50 kDa | 0 | 3 | 2 | 5 |
| Catenin alpha-1 OS=Homo sapiens OX=9606 GN=CTNNA1 PE=1 SV=1 | P35221 (+1) | CTNNA1 | 100 kDa | 1 | 4 | 0 | 5 |

| | | | | | | | |
|---|---|---|---|---|---|---|---|
| Collagen alpha-1(XVIII) chain OS=Homo sapiens OX=9606 GN=COL18A1 PE=1 SV=5 | P39060 (+2) | COL18A1 | 178 kDa | 0 | 1 | 4 | 5 |
| Microtubule-actin cross-linking factor 1, isoforms 1/2/3/5 OS=Homo sapiens OX=9606 GN=MACF1 PE=1 SV=4 | Q9UPN3 | MACF1 | 838 kDa | 0 | 2 | 3 | 5 |
| Isoform 4 of A-kinase anchor protein 2 OS=Homo sapiens OX=9606 GN=AKAP2 | Q9Y2D5-6 | AKAP2 | 121 kDa | 1 | 2 | 2 | 5 |
| Leucine--tRNA ligase, cytoplasmic OS=Homo sapiens OX=9606 GN=LARS1 PE=1 SV=2 | Q9P2J5 (+1) | LARS1 | 134 kDa | 1 | 2 | 2 | 5 |
| Histone H3.1 OS=Homo sapiens OX=9606 GN=H3C1 PE=1 SV=2 | P68431 (+1) | H3C1 | 15 kDa | 2 | 1 | 2 | 5 |
| PDZ and LIM domain protein 7 OS=Homo sapiens OX=9606 GN=PDLIM7 PE=1 SV=1 | Q9NR12 | PDLIM7 | 50 kDa | 2 | 2 | 1 | 5 |
| 40S ribosomal protein S3a OS=Homo sapiens OX=9606 GN=RPS3A PE=1 SV=2 | P61247 | RPS3A | 30 kDa | 1 | 2 | 2 | 5 |
| Protein disulfide-isomerase A4 OS=Homo sapiens OX=9606 GN=PDIA4 PE=1 SV=2 | P13667 | PDIA4 | 73 kDa | 2 | 0 | 3 | 5 |
| Heterogeneous nuclear ribonucleoprotein U-like protein 2 OS=Homo sapiens OX=9606 GN=HNRNPUL2 PE=1 SV=1 | Q1KMD3 | HNRNPUL2 | 85 kDa | 2 | 2 | 1 | 5 |
| Extended synaptotagmin-1 OS=Homo sapiens OX=9606 GN=ESYT1 PE=1 SV=1 | Q9BSJ8 (+1) | ESYT1 | 123 kDa | 0 | 3 | 2 | 5 |
| Isoform Beta-4B of Integrin beta-4 OS=Homo sapiens OX=9606 GN=ITGB4 | P16144-3 | ITGB4 | 201 kDa | 1 | 0 | 4 | 5 |
| Cysteine-rich protein 2 OS=Homo sapiens OX=9606 GN=CRIP2 PE=1 SV=1 | P52943 (+1) | CRIP2 | 22 kDa | 1 | 2 | 2 | 5 |
| Insulin-like growth factor 2 mRNA-binding protein 3 OS=Homo sapiens OX=9606 GN=IGF2BP3 PE=1 SV=2 | O00425 | IGF2BP3 | 64 kDa | 0 | 3 | 2 | 5 |
| Triosephosphate isomerase OS=Homo sapiens OX=9606 GN=TPI1 PE=1 SV=4 | P60174 (+1) | TPI1 | 27 kDa | 2 | 1 | 2 | 5 |
| Drebrin OS=Homo sapiens OX=9606 GN=DBN1 PE=1 SV=4 | Q16643 (+2) | DBN1 | 71 kDa | 1 | 2 | 2 | 5 |
| Isoform LCRMP-4 of Dihydropyrimidinase-related protein 3 OS=Homo sapiens OX=9606 GN=DPYSL3 | Q14195-2 | DPYSL3 | 74 kDa | 1 | 0 | 4 | 5 |
| S-adenosylmethionine synthase isoform type-2 OS=Homo sapiens OX=9606 GN=MAT2A PE=1 SV=1 | P31153 | MAT2A | 44 kDa | 2 | 2 | 1 | 5 |

| | | | | | | | |
|---|---|---|---|---|---|---|---|
| Calponin-2 OS=Homo sapiens OX=9606 GN=CNN2 PE=1 SV=4 | Q99439 | CNN2 | 34 kDa | 1 | 2 | 2 | 5 |
| Transportin-1 OS=Homo sapiens OX=9606 GN=TNPO1 PE=1 SV=2 | Q92973 (+2) | TNPO1 | 102 kDa | 1 | 2 | 2 | 5 |
| Protein phosphatase 1F OS=Homo sapiens OX=9606 GN=PPM1F PE=1 SV=3 | P49593 | PPM1F | 50 kDa | 2 | 0 | 3 | 5 |
| ATP-dependent 6-phosphofructokinase, liver type OS=Homo sapiens OX=9606 GN=PFKL PE=1 SV=6 | P17858 | PFKL | 85 kDa | 1 | 2 | 2 | 5 |
| Ras-related C3 botulinum toxin substrate 1 OS=Homo sapiens OX=9606 GN=RAC1 PE=1 SV=1 | P63000 (+1) | RAC1 | 21 kDa | 1 | 2 | 2 | 5 |
| Protein phosphatase 1 regulatory subunit 12A OS=Homo sapiens OX=9606 GN=PPP1R12A PE=1 SV=1 | O14974 (+4) | PPP1R12A | 115 kDa | 2 | 3 | 0 | 5 |
| Mitogen-activated protein kinase 1 OS=Homo sapiens OX=9606 GN=MAPK1 PE=1 SV=3 | P28482 | MAPK1 | 41 kDa | 0 | 3 | 2 | 5 |
| Liprin-beta-1 OS=Homo sapiens OX=9606 GN=PPFIBP1 PE=1 SV=2 | Q86W92 (+3) | PPFIBP1 | 114 kDa | 1 | 2 | 2 | 5 |
| Mitogen-activated protein kinase 3 OS=Homo sapiens OX=9606 GN=MAPK3 PE=1 SV=4 | P27361 | MAPK3 | 43 kDa | 2 | 2 | 1 | 5 |
| Palladin OS=Homo sapiens OX=9606 GN=PALLD PE=1 SV=3 | Q8WX93 (+4) | PALLD | 151 kDa | 2 | 2 | 1 | 5 |
| Glutamate dehydrogenase 1, mitochondrial OS=Homo sapiens OX=9606 GN=GLUD1 PE=1 SV=2 | P00367 (+1) | GLUD1 | 61 kDa | 1 | 2 | 2 | 5 |
| F-actin-capping protein subunit beta OS=Homo sapiens OX=9606 GN=CAPZB PE=1 SV=4 | P47756 (+1) | CAPZB | 31 kDa | 1 | 2 | 2 | 5 |
| Utrophin OS=Homo sapiens OX=9606 GN=UTRN PE=1 SV=2 | P46939 | UTRN | 394 kDa | 0 | 3 | 2 | 5 |
| Alanine--tRNA ligase, cytoplasmic OS=Homo sapiens OX=9606 GN=AARS1 PE=1 SV=2 | P49588 | AARS1 | 107 kDa | 1 | 1 | 3 | 5 |
| Eukaryotic translation initiation factor 3 subunit M OS=Homo sapiens OX=9606 GN=EIF3M PE=1 SV=1 | Q7L2H7 | EIF3M | 43 kDa | 3 | 1 | 1 | 5 |
| 14-3-3 protein gamma OS=Homo sapiens OX=9606 GN=YWHAG PE=1 SV=2 | P61981 | YWHAG | 28 kDa | 1 | 1 | 3 | 5 |
| Isoform 5 of RNA-binding protein 14 OS=Homo sapiens OX=9606 GN=RBM14 | Q96PK6-5 (+1) | RBM14 | 37 kDa | 1 | 2 | 2 | 5 |
| Nidogen-1 OS=Homo sapiens OX=9606 GN=NID1 PE=1 SV=3 | P14543 (+1) | NID1 | 136 kDa | 1 | 1 | 3 | 5 |

| | | | | | | | |
|---|---|---|---|---|---|---|---|
| Pre-mRNA-processing-splicing factor 8 OS=Homo sapiens OX=9606 GN=PRPF8 PE=1 SV=2 | Q6P2Q9 | PRPF8 | 274 kDa | 2 | 1 | 2 | 5 |
| Eukaryotic translation initiation factor 4 gamma 1 OS=Homo sapiens OX=9606 GN=EIF4G1 PE=1 SV=4 | Q04637 (+4) | EIF4G1 | 175 kDa | 0 | 2 | 3 | 5 |
| Serine--tRNA ligase, cytoplasmic OS=Homo sapiens OX=9606 GN=SARS1 PE=1 SV=3 | P49591 | SARS1 | 59 kDa | 2 | 1 | 2 | 5 |
| Puromycin-sensitive aminopeptidase OS=Homo sapiens OX=9606 GN=NPEPPS PE=1 SV=2 | P55786 | NPEPPS | 103 kDa | 2 | 1 | 2 | 5 |
| Adenosylhomocysteinase OS=Homo sapiens OX=9606 GN=AHCY PE=1 SV=4 | P23526 | AHCY | 48 kDa | 1 | 2 | 2 | 5 |
| E3 SUMO-protein ligase RanBP2 OS=Homo sapiens OX=9606 GN=RANBP2 PE=1 SV=2 | P49792 | RANBP2 | 358 kDa | 0 | 2 | 3 | 5 |
| Polyadenylate-binding protein 1 OS=Homo sapiens OX=9606 GN=PABPC1 PE=1 SV=2 | P11940 (+1) | PABPC1 | 71 kDa | 2 | 1 | 1 | 4 |
| Serpin H1 OS=Homo sapiens OX=9606 GN=SERPINH1 PE=1 SV=2 | P50454 | SERPINH1 | 46 kDa | 2 | 1 | 1 | 4 |
| Isoform 3 of Heterogeneous nuclear ribonucleoprotein D0 OS=Homo sapiens OX=9606 GN=HNRNPD | Q14103-3 (+1) | HNRNPD | 33 kDa | 1 | 2 | 1 | 4 |
| Proliferation-associated protein 2G4 OS=Homo sapiens OX=9606 GN=PA2G4 PE=1 SV=3 | Q9UQ80 | PA2G4 | 44 kDa | 1 | 1 | 2 | 4 |
| Protein unc-45 homolog A OS=Homo sapiens OX=9606 GN=UNC45A PE=1 SV=1 | Q9H3U1 (+1) | UNC45A | 103 kDa | 0 | 1 | 3 | 4 |
| T-complex protein 1 subunit eta OS=Homo sapiens OX=9606 GN=CCT7 PE=1 SV=2 | Q99832 | CCT7 | 59 kDa | 1 | 0 | 3 | 4 |
| Exportin-2 OS=Homo sapiens OX=9606 GN=CSE1L PE=1 SV=3 | P55060 (+2) | CSE1L | 110 kDa | 0 | 2 | 2 | 4 |
| KH domain-containing, RNA-binding, signal transduction-associated protein 1 OS=Homo sapiens OX=9606 GN=KHDRBS1 PE=1 SV=1 | Q07666 | KHDRBS1 | 48 kDa | 2 | 1 | 1 | 4 |
| Far upstream element-binding protein 3 OS=Homo sapiens OX=9606 GN=FUBP3 PE=1 SV=2 | Q96I24 | FUBP3 | 62 kDa | 0 | 2 | 2 | 4 |
| 5'-3' exoribonuclease 2 OS=Homo sapiens OX=9606 GN=XRN2 PE=1 SV=1 | Q9H0D6 (+1) | XRN2 | 109 kDa | 1 | 2 | 1 | 4 |

| | | | | | | | |
|---|---|---|---|---|---|---|---|
| Isoform 2 of Hypoxia up-regulated protein 1 OS=Homo sapiens OX=9606 GN=HYOU1 | Q9Y4L1-2 | HYOU1 | 63 kDa | 0 | 2 | 2 | 4 |
| Splicing factor 3A subunit 3 OS=Homo sapiens OX=9606 GN=SF3A3 PE=1 SV=1 | Q12874 | SF3A3 | 59 kDa | 0 | 3 | 1 | 4 |
| RuvB-like 2 OS=Homo sapiens OX=9606 GN=RUVBL2 PE=1 SV=3 | Q9Y230 | RUVBL2 | 51 kDa | 1 | 1 | 2 | 4 |
| Ezrin OS=Homo sapiens OX=9606 GN=EZR PE=1 SV=4 | P15311 | EZR | 69 kDa | 2 | 1 | 1 | 4 |
| Far upstream element-binding protein 1 OS=Homo sapiens OX=9606 GN=FUBP1 PE=1 SV=3 | Q96AE4 (+1) | FUBP1 | 68 kDa | 2 | 0 | 2 | 4 |
| RNA-binding protein FUS OS=Homo sapiens OX=9606 GN=FUS PE=1 SV=1 | P35637 (+1) | FUS | 53 kDa | 1 | 2 | 1 | 4 |
| Asparagine--tRNA ligase, cytoplasmic OS=Homo sapiens OX=9606 GN=NARS1 PE=1 SV=1 | O43776 | NARS1 | 63 kDa | 1 | 1 | 2 | 4 |
| Isoform 2 of Gelsolin OS=Homo sapiens OX=9606 GN=GSN | P06396-2 | GSN | 81 kDa | 1 | 2 | 1 | 4 |
| Glutamine--tRNA ligase OS=Homo sapiens OX=9606 GN=QARS1 PE=1 SV=1 | P47897 (+1) | QARS1 | 88 kDa | 2 | 0 | 2 | 4 |
| 1-phosphatidylinositol 4,5-bisphosphate phosphodiesterase beta-3 OS=Homo sapiens OX=9606 GN=PLCB3 PE=1 SV=2 | Q01970 | PLCB3 | 139 kDa | 1 | 2 | 1 | 4 |
| 40S ribosomal protein S11 OS=Homo sapiens OX=9606 GN=RPS11 PE=1 SV=3 | P62280 | RPS11 | 18 kDa | 1 | 1 | 2 | 4 |
| Phenylalanine--tRNA ligase alpha subunit OS=Homo sapiens OX=9606 GN=FARSA PE=1 SV=3 | Q9Y285 (+1) | FARSA | 58 kDa | 2 | 1 | 1 | 4 |
| Annexin A7 OS=Homo sapiens OX=9606 GN=ANXA7 PE=1 SV=3 | P20073 (+1) | ANXA7 | 53 kDa | 1 | 1 | 2 | 4 |
| Dedicator of cytokinesis protein 6 OS=Homo sapiens OX=9606 GN=DOCK6 PE=1 SV=3 | Q96HP0 | DOCK6 | 230 kDa | 1 | 1 | 2 | 4 |
| Spermidine synthase OS=Homo sapiens OX=9606 GN=SRM PE=1 SV=1 | P19623 | SRM | 34 kDa | 1 | 1 | 2 | 4 |
| Fragile X mental retardation syndrome-related protein 1 OS=Homo sapiens OX=9606 GN=FXR1 PE=1 SV=3 | P51114 (+1) | FXR1 | 70 kDa | 2 | 0 | 2 | 4 |
| Tight junction protein ZO-2 OS=Homo sapiens OX=9606 GN=TJP2 PE=1 SV=2 | Q9UDY2 (+3) | TJP2 | 134 kDa | 0 | 3 | 1 | 4 |

| | | | | | | | |
|---|---|---|---|---|---|---|---|
| Ran GTPase-activating protein 1 OS=Homo sapiens OX=9606 GN=RANGAP1 PE=1 SV=1 | P46060 | RANGAP1 | 64 kDa | 1 | 1 | 2 | 4 |
| Acylamino-acid-releasing enzyme OS=Homo sapiens OX=9606 GN=APEH PE=1 SV=4 | P13798 | APEH | 81 kDa | 0 | 2 | 2 | 4 |
| Caveolae-associated protein 1 OS=Homo sapiens OX=9606 GN=CAVIN1 PE=1 SV=1 | Q6NZI2 | CAVIN1 | 43 kDa | 3 | 1 | 0 | 4 |
| Procollagen galactosyltransferase 1 OS=Homo sapiens OX=9606 GN=COLGALT1 PE=1 SV=1 | Q8NBJ5 | COLGALT1 | 72 kDa | 1 | 2 | 1 | 4 |
| Eukaryotic translation elongation factor 1 epsilon-1 OS=Homo sapiens OX=9606 GN=EEF1E1 PE=1 SV=1 | O43324 (+1) | EEF1E1 | 20 kDa | 2 | 1 | 1 | 4 |
| 60S ribosomal protein L3 OS=Homo sapiens OX=9606 GN=RPL3 PE=1 SV=2 | P39023 | RPL3 | 46 kDa | 1 | 1 | 2 | 4 |
| Rho GDP-dissociation inhibitor 1 OS=Homo sapiens OX=9606 GN=ARHGDIA PE=1 SV=3 | P52565 | ARHGDIA | 23 kDa | 1 | 0 | 3 | 4 |
| Cysteine and glycine-rich protein 1 OS=Homo sapiens OX=9606 GN=CSRP1 PE=1 SV=3 | P21291 | CSRP1 | 21 kDa | 2 | 1 | 1 | 4 |
| E3 ubiquitin-protein ligase RNF213 OS=Homo sapiens OX=9606 GN=RNF213 PE=1 SV=3 | Q63HN8 (+1) | RNF213 | 591 kDa | 1 | 2 | 1 | 4 |
| H/ACA ribonucleoprotein complex subunit DKC1 OS=Homo sapiens OX=9606 GN=DKC1 PE=1 SV=3 | O60832 (+1) | DKC1 | 58 kDa | 1 | 0 | 3 | 4 |
| Centrosomal protein of 170 kDa OS=Homo sapiens OX=9606 GN=CEP170 PE=1 SV=1 | Q5SW79 (+2) | CEP170 | 175 kDa | 1 | 0 | 3 | 4 |
| Endophilin-A2 OS=Homo sapiens OX=9606 GN=SH3GL1 PE=1 SV=1 | Q99961 | SH3GL1 | 41 kDa | 1 | 2 | 1 | 4 |
| Polyadenylate-binding protein 2 OS=Homo sapiens OX=9606 GN=PABPN1 PE=1 SV=3 | Q86U42 (+1) | PABPN1 | 33 kDa | 2 | 1 | 1 | 4 |
| Threonine--tRNA ligase 1, cytoplasmic OS=Homo sapiens OX=9606 GN=TARS1 PE=1 SV=3 | P26639 | TARS1 | 83 kDa | 2 | 1 | 1 | 4 |
| Elongation factor Tu, mitochondrial OS=Homo sapiens OX=9606 GN=TUFM PE=1 SV=2 | P49411 | TUFM | 50 kDa | 1 | 1 | 2 | 4 |
| GTP-binding nuclear protein Ran OS=Homo sapiens OX=9606 GN=RAN PE=1 SV=3 | P62826 | RAN | 24 kDa | 1 | 0 | 3 | 4 |

| | | | | | | | |
|---|---|---|---|---|---|---|---|
| 14-3-3 protein eta OS=Homo sapiens OX=9606 GN=YWHAH PE=1 SV=4 | Q04917 | YWHAH | 28 kDa | 1 | 2 | 1 | 4 |
| TAR DNA-binding protein 43 OS=Homo sapiens OX=9606 GN=TARDBP PE=1 SV=1 | Q13148 (+1) | TARDBP | 45 kDa | 1 | 2 | 1 | 4 |
| FACT complex subunit SPT16 OS=Homo sapiens OX=9606 GN=SUPT16H PE=1 SV=1 | Q9Y5B9 | SUPT16H | 120 kDa | 1 | 1 | 2 | 4 |
| Isoform 2 of HLA class I histocompatibility antigen, A alpha chain OS=Homo sapiens OX=9606 GN=HLA-A | P04439-2 | HLA-A | 41 kDa | 1 | 0 | 3 | 4 |
| Stress-induced-phosphoprotein 1 OS=Homo sapiens OX=9606 GN=STIP1 PE=1 SV=1 | P31948 | STIP1 | 63 kDa | 0 | 2 | 2 | 4 |
| Beta-2-syntrophin OS=Homo sapiens OX=9606 GN=SNTB2 PE=1 SV=1 | Q13425 (+1) | SNTB2 | 58 kDa | 1 | 2 | 1 | 4 |
| Unconventional myosin-Va OS=Homo sapiens OX=9606 GN=MYO5A PE=1 SV=2 | Q9Y4I1 (+2) | MYO5A | 215 kDa | 1 | 2 | 1 | 4 |
| Importin-7 OS=Homo sapiens OX=9606 GN=IPO7 PE=1 SV=1 | O95373 | IPO7 | 120 kDa | 0 | 2 | 2 | 4 |
| Zinc finger CCCH-type antiviral protein 1 OS=Homo sapiens OX=9606 GN=ZC3HAV1 PE=1 SV=3 | Q7Z2W4 | ZC3HAV1 | 101 kDa | 1 | 1 | 2 | 4 |
| Actin-related protein 2/3 complex subunit 1B OS=Homo sapiens OX=9606 GN=ARPC1B PE=1 SV=3 | O15143 | ARPC1B | 41 kDa | 2 | 0 | 2 | 4 |
| Kinectin OS=Homo sapiens OX=9606 GN=KTN1 PE=1 SV=1 | Q86UP2 (+3) | KTN1 | 156 kDa | 2 | 1 | 1 | 4 |
| Abl interactor 1 OS=Homo sapiens OX=9606 GN=ABI1 PE=1 SV=4 | Q8IZP0 (+11) | ABI1 | 55 kDa | 1 | 1 | 2 | 4 |
| Atlastin-3 OS=Homo sapiens OX=9606 GN=ATL3 PE=1 SV=1 | Q6DD88 | ATL3 | 61 kDa | 0 | 2 | 2 | 4 |
| Adenine phosphoribosyltransferase OS=Homo sapiens OX=9606 GN=APRT PE=1 SV=2 | P07741 | APRT | 20 kDa | 2 | 1 | 1 | 4 |
| Ataxin-2 OS=Homo sapiens OX=9606 GN=ATXN2 PE=1 SV=2 | Q99700 (+3) | ATXN2 | 140 kDa | 2 | 1 | 1 | 4 |
| Splicing factor 3B subunit 3 OS=Homo sapiens OX=9606 GN=SF3B3 PE=1 SV=4 | Q15393 | SF3B3 | 136 kDa | 1 | 1 | 1 | 3 |
| Poly(rC)-binding protein 1 OS=Homo sapiens OX=9606 GN=PCBP1 PE=1 SV=2 | Q15365 | PCBP1 | 37 kDa | 0 | 2 | 1 | 3 |
| U5 small nuclear ribonucleoprotein 200 kDa helicase OS=Homo sapiens OX=9606 GN=SNRNP200 PE=1 SV=2 | O75643 | SNRNP200 | 245 kDa | 1 | 1 | 1 | 3 |

| | | | | | | | |
|---|---|---|---|---|---|---|---|
| Leucine-rich PPR motif-containing protein, mitochondrial OS=Homo sapiens OX=9606 GN=LRPPRC PE=1 SV=3 | P42704 | LRPPRC | 158 kDa | 1 | 1 | 1 | 3 |
| Vacuolar protein sorting-associated protein 35 OS=Homo sapiens OX=9606 GN=VPS35 PE=1 SV=2 | Q96QK1 | VPS35 | 92 kDa | 1 | 1 | 1 | 3 |
| Lysosomal Pro-X carboxypeptidase OS=Homo sapiens OX=9606 GN=PRCP PE=1 SV=1 | P42785 (+1) | PRCP | 56 kDa | 0 | 1 | 2 | 3 |
| Cullin-associated NEDD8-dissociated protein 1 OS=Homo sapiens OX=9606 GN=CAND1 PE=1 SV=2 | Q86VP6 | CAND1 | 136 kDa | 1 | 1 | 1 | 3 |
| Guanine nucleotide-binding protein G(I)/G(S)/G(T) subunit beta-1 OS=Homo sapiens OX=9606 GN=GNB1 PE=1 SV=3 | P62873 (+1) | GNB1 | 37 kDa | 1 | 1 | 1 | 3 |
| Protein PML OS=Homo sapiens OX=9606 GN=PML PE=1 SV=3 | P29590 (+1) | PML | 98 kDa | 1 | 1 | 1 | 3 |
| Multimerin-2 OS=Homo sapiens OX=9606 GN=MMRN2 PE=1 SV=2 | Q9H8L6 | MMRN2 | 104 kDa | 0 | 1 | 2 | 3 |
| Eukaryotic translation initiation factor 4 gamma 2 OS=Homo sapiens OX=9606 GN=EIF4G2 PE=1 SV=1 | P78344 | EIF4G2 | 102 kDa | 1 | 1 | 1 | 3 |
| Isoform 6 of Thioredoxin reductase 1, cytoplasmic OS=Homo sapiens OX=9606 GN=TXNRD1 | Q16881-6 | TXNRD1 | 67 kDa | 1 | 0 | 2 | 3 |
| Dynamin-2 OS=Homo sapiens OX=9606 GN=DNM2 PE=1 SV=2 | P50570 (+2) | DNM2 | 98 kDa | 0 | 2 | 1 | 3 |
| Heterogeneous nuclear ribonucleoprotein A3 OS=Homo sapiens OX=9606 GN=HNRNPA3 PE=1 SV=2 | P51991 | HNRNPA3 | 40 kDa | 1 | 1 | 1 | 3 |
| DNA damage-binding protein 1 OS=Homo sapiens OX=9606 GN=DDB1 PE=1 SV=1 | Q16531 | DDB1 | 127 kDa | 1 | 0 | 2 | 3 |
| Tropomyosin alpha-4 chain OS=Homo sapiens OX=9606 GN=TPM4 PE=1 SV=3 | P67936 | TPM4 | 29 kDa | 0 | 2 | 1 | 3 |
| Importin-5 OS=Homo sapiens OX=9606 GN=IPO5 PE=1 SV=4 | O00410 | IPO5 | 124 kDa | 1 | 0 | 2 | 3 |
| 182 kDa tankyrase-1-binding protein OS=Homo sapiens OX=9606 GN=TNKS1BP1 PE=1 SV=4 | Q9C0C2 | TNKS1BP1 | 182 kDa | 1 | 2 | 0 | 3 |
| Toll-interacting protein OS=Homo sapiens OX=9606 GN=TOLLIP PE=1 SV=1 | Q9H0E2 | TOLLIP | 30 kDa | 1 | 0 | 2 | 3 |
| Xaa-Pro aminopeptidase 1 OS=Homo sapiens OX=9606 GN=XPNPEP1 PE=1 SV=3 | Q9NQW7 | XPNPEP1 | 70 kDa | 0 | 1 | 2 | 3 |

| | | | | | | | |
|---|---|---|---|---|---|---|---|
| Integrin-linked protein kinase OS=Homo sapiens OX=9606 GN=ILK PE=1 SV=2 | Q13418 (+1) | ILK | 51 kDa | 0 | 1 | 2 | 3 |
| Caldesmon OS=Homo sapiens OX=9606 GN=CALD1 PE=1 SV=3 | Q05682 | CALD1 | 93 kDa | 1 | 2 | 0 | 3 |
| Rho-related GTP-binding protein RhoC OS=Homo sapiens OX=9606 GN=RHOC PE=1 SV=1 | P08134 (+1) | RHOC | 22 kDa | 0 | 1 | 2 | 3 |
| Serine/arginine-rich splicing factor 6 OS=Homo sapiens OX=9606 GN=SRSF6 PE=1 SV=2 | Q13247 (+1) | SRSF6 | 40 kDa | 1 | 1 | 1 | 3 |
| ATP-binding cassette sub-family E member 1 OS=Homo sapiens OX=9606 GN=ABCE1 PE=1 SV=1 | P61221 | ABCE1 | 67 kDa | 1 | 0 | 2 | 3 |
| Leucine zipper protein 1 OS=Homo sapiens OX=9606 GN=LUZP1 PE=1 SV=2 | Q86V48 (+2) | LUZP1 | 120 kDa | 0 | 1 | 2 | 3 |
| Ras-related protein Rap-1b-like protein OS=Homo sapiens OX=9606 PE=2 SV=1 | A6NIZ1 (+4) | RP1BL | 21 kDa | 1 | 0 | 2 | 3 |
| Twinfilin-1 OS=Homo sapiens OX=9606 GN=TWF1 PE=1 SV=3 | Q12792 (+1) | TWF1 | 40 kDa | 1 | 1 | 1 | 3 |
| 26S proteasome non-ATPase regulatory subunit 12 OS=Homo sapiens OX=9606 GN=PSMD12 PE=1 SV=3 | O00232 | PSMD12 | 53 kDa | 1 | 1 | 1 | 3 |
| Pre-mRNA-processing factor 19 OS=Homo sapiens OX=9606 GN=PRPF19 PE=1 SV=1 | Q9UMS4 | PRPF19 | 55 kDa | 1 | 1 | 1 | 3 |
| Interleukin enhancer-binding factor 3 OS=Homo sapiens OX=9606 GN=ILF3 PE=1 SV=3 | Q12906 (+6) | ILF3 | 95 kDa | 0 | 2 | 1 | 3 |
| Guanine nucleotide-binding protein G(I)/G(S)/G(T) subunit beta-2 OS=Homo sapiens OX=9606 GN=GNB2 PE=1 SV=3 | P62879 | GNB2 | 37 kDa | 1 | 1 | 1 | 3 |
| Y-box-binding protein 1 OS=Homo sapiens OX=9606 GN=YBX1 PE=1 SV=3 | P67809 | YBX1 | 36 kDa | 1 | 0 | 2 | 3 |
| RNA-splicing ligase RtcB homolog OS=Homo sapiens OX=9606 GN=RTCB PE=1 SV=1 | Q9Y3I0 | RTCB | 55 kDa | 0 | 0 | 3 | 3 |
| Exosome complex component RRP40 OS=Homo sapiens OX=9606 GN=EXOSC3 PE=1 SV=3 | Q9NQT5 (+1) | EXOSC3 | 30 kDa | 1 | 0 | 2 | 3 |
| Calpain small subunit 1 OS=Homo sapiens OX=9606 GN=CAPNS1 PE=1 SV=1 | P04632 | CAPNS1 | 28 kDa | 1 | 0 | 2 | 3 |

| | | | | | | | |
|---|---|---|---|---|---|---|---|
| 26S proteasome regulatory subunit 7 OS=Homo sapiens OX=9606 GN=PSMC2 PE=1 SV=3 | P35998 | PSMC2 | 49 kDa | 1 | 0 | 2 | 3 |
| RNA-binding protein 14 OS=Homo sapiens OX=9606 GN=RBM14 PE=1 SV=2 | Q96PK6 | RBM14 | 69 kDa | 1 | 2 | 0 | 3 |
| Adenylyl cyclase-associated protein 1 OS=Homo sapiens OX=9606 GN=CAP1 PE=1 SV=5 | Q01518 (+1) | CAP1 | 52 kDa | 0 | 1 | 2 | 3 |
| 60S ribosomal protein L7a OS=Homo sapiens OX=9606 GN=RPL7A PE=1 SV=2 | P62424 | RPL7A | 30 kDa | 1 | 1 | 1 | 3 |
| Carnitine O-palmitoyltransferase 1, liver isoform OS=Homo sapiens OX=9606 GN=CPT1A PE=1 SV=2 | P50416 (+1) | CPT1A | 88 kDa | 1 | 1 | 1 | 3 |
| Stathmin OS=Homo sapiens OX=9606 GN=STMN1 PE=1 SV=3 | P16949 (+1) | STMN1 | 17 kDa | 0 | 2 | 1 | 3 |
| Inosine-5'-monophosphate dehydrogenase 2 OS=Homo sapiens OX=9606 GN=IMPDH2 PE=1 SV=2 | P12268 | IMPDH2 | 56 kDa | 0 | 2 | 1 | 3 |
| 14-3-3 protein epsilon OS=Homo sapiens OX=9606 GN=YWHAE PE=1 SV=1 | P62258 | YWHAE | 29 kDa | 0 | 2 | 1 | 3 |
| Signal recognition particle subunit SRP68 OS=Homo sapiens OX=9606 GN=SRP68 PE=1 SV=2 | Q9UHB9 (+1) | SRP68 | 71 kDa | 2 | 0 | 1 | 3 |
| Isoform 2 of Macrophage-capping protein OS=Homo sapiens OX=9606 GN=CAPG | P40121-2 | CAPG | 37 kDa | 1 | 0 | 2 | 3 |
| Hsp90 co-chaperone Cdc37 OS=Homo sapiens OX=9606 GN=CDC37 PE=1 SV=1 | Q16543 | CDC37 | 44 kDa | 0 | 3 | 0 | 3 |
| Core histone macro-H2A.1 OS=Homo sapiens OX=9606 GN=MACROH2A1 PE=1 SV=4 | O75367 (+2) | MACROH2A1 | 40 kDa | 1 | 0 | 2 | 3 |
| Dynein axonemal assembly factor 5 OS=Homo sapiens OX=9606 GN=DNAAF5 PE=1 SV=4 | Q86Y56 (+1) | DNAAF5 | 94 kDa | 2 | 0 | 1 | 3 |
| Dihydrolipoyllysine-residue acetyltransferase component of pyruvate dehydrogenase complex, mitochondrial OS=Homo sapiens OX=9606 GN=DLAT PE=1 SV=3 | P10515 | DLAT | 69 kDa | 1 | 0 | 2 | 3 |
| 40S ribosomal protein S3 OS=Homo sapiens OX=9606 GN=RPS3 PE=1 SV=2 | P23396 | RPS3 | 27 kDa | 0 | 1 | 2 | 3 |
| Histone H2B type 1-O OS=Homo sapiens OX=9606 GN=H2BC17 PE=1 SV=3 | P23527 (+10) | H2BC17 | 14 kDa | 1 | 0 | 2 | 3 |

| | | | | | | | |
|---|---|---|---|---|---|---|---|
| Isoform 3 of LIM domain only protein 7 OS=Homo sapiens OX=9606 GN=LMO7 | Q8WWI1-3 | LMO7 | 154 kDa | 0 | 2 | 1 | 3 |
| Coiled-coil domain-containing protein 22 OS=Homo sapiens OX=9606 GN=CCDC22 PE=1 SV=1 | O60826 | CCDC22 | 71 kDa | 0 | 1 | 2 | 3 |
| GRB10-interacting GYF protein 2 OS=Homo sapiens OX=9606 GN=GIGYF2 PE=1 SV=1 | Q6Y7W6 (+3) | GIGYF2 | 150 kDa | 1 | 2 | 0 | 3 |
| Glutamine synthetase OS=Homo sapiens OX=9606 GN=GLUL PE=1 SV=4 | P15104 | GLUL | 42 kDa | 2 | 0 | 1 | 3 |
| Keratin, type I cytoskeletal 9 OS=Homo sapiens OX=9606 GN=KRT9 PE=1 SV=3 | P35527 (+1) | KRT9 | 62 kDa | 0 | 1 | 1 | 2 |
| Arginine--tRNA ligase, cytoplasmic OS=Homo sapiens OX=9606 GN=RARS1 PE=1 SV=2 | P54136 | RARS1 | 75 kDa | 0 | 1 | 1 | 2 |
| Hexokinase-1 OS=Homo sapiens OX=9606 GN=HK1 PE=1 SV=3 | P19367 (+3) | HK1 | 102 kDa | 1 | 1 | 0 | 2 |
| Adenylosuccinate synthetase isozyme 2 OS=Homo sapiens OX=9606 GN=ADSS2 PE=1 SV=3 | P30520 | ADSS2 | 50 kDa | 1 | 0 | 1 | 2 |
| Serine/threonine-protein phosphatase 2A 65 kDa regulatory subunit A alpha isoform OS=Homo sapiens OX=9606 GN=PPP2R1A PE=1 SV=4 | P30153 | PPP2R1A | 65 kDa | 0 | 1 | 1 | 2 |
| PDZ and LIM domain protein 5 OS=Homo sapiens OX=9606 GN=PDLIM5 PE=1 SV=5 | Q96HC4 | PDLIM5 | 64 kDa | 1 | 1 | 0 | 2 |
| Calpain-1 catalytic subunit OS=Homo sapiens OX=9606 GN=CAPN1 PE=1 SV=1 | P07384 | CAPN1 | 82 kDa | 0 | 1 | 1 | 2 |
| Isoform 3 of Septin-2 OS=Homo sapiens OX=9606 GN=SEPTIN2 | Q15019-3 | SEPTIN2 | 43 kDa | 0 | 1 | 1 | 2 |
| Glucosidase 2 subunit beta OS=Homo sapiens OX=9606 GN=PRKCSH PE=1 SV=2 | P14314 (+1) | PRKCSH | 59 kDa | 0 | 1 | 1 | 2 |
| N-acetylglucosamine-6-sulfatase OS=Homo sapiens OX=9606 GN=GNS PE=1 SV=3 | P15586 (+1) | GNS | 62 kDa | 1 | 0 | 1 | 2 |
| Myosin-10 OS=Homo sapiens OX=9606 GN=MYH10 PE=1 SV=3 | P35580 (+4) | MYH10 | 229 kDa | 1 | 0 | 1 | 2 |
| Heterogeneous nuclear ribonucleoprotein Q OS=Homo sapiens OX=9606 GN=SYNCRIP PE=1 SV=2 | O60506 | SYNCRIP | 70 kDa | 0 | 1 | 1 | 2 |

| | | | | | | | |
|---|---|---|---|---|---|---|---|
| Endoglin OS=Homo sapiens OX=9606 GN=ENG PE=1 SV=2 | P17813 (+1) | ENG | 71 kDa | 0 | 1 | 1 | 2 |
| Heterogeneous nuclear ribonucleoprotein U-like protein 1 OS=Homo sapiens OX=9606 GN=HNRNPUL1 PE=1 SV=2 | Q9BUJ2 (+2) | HNRNPUL1 | 96 kDa | 1 | 0 | 1 | 2 |
| 3-mercaptopyruvate sulfurtransferase OS=Homo sapiens OX=9606 GN=MPST PE=1 SV=3 | P25325 (+1) | MPST | 33 kDa | 1 | 1 | 0 | 2 |
| Tripeptidyl-peptidase 1 OS=Homo sapiens OX=9606 GN=TPP1 PE=1 SV=2 | O14773 (+1) | TPP1 | 61 kDa | 0 | 1 | 1 | 2 |
| Regulator of nonsense transcripts 1 OS=Homo sapiens OX=9606 GN=UPF1 PE=1 SV=2 | Q92900 | UPF1 | 124 kDa | 0 | 2 | 0 | 2 |
| Keratin, type II cytoskeletal 2 epidermal OS=Homo sapiens OX=9606 GN=KRT2 PE=1 SV=2 | P35908 (+2) | KRT2 | 65 kDa | 1 | 1 | 0 | 2 |
| Tyrosine--tRNA ligase, cytoplasmic OS=Homo sapiens OX=9606 GN=YARS1 PE=1 SV=4 | P54577 | YARS1 | 59 kDa | 1 | 0 | 1 | 2 |
| Trifunctional purine biosynthetic protein adenosine-3 OS=Homo sapiens OX=9606 GN=GART PE=1 SV=1 | P22102 | GART | 108 kDa | 1 | 1 | 0 | 2 |
| Isoform Non-brain of Clathrin light chain A OS=Homo sapiens OX=9606 GN=CLTA | P09496-2 (+1) | CLTA | 24 kDa | 0 | 1 | 1 | 2 |
| Multifunctional procollagen lysine hydroxylase and glycosyltransferase LH3 OS=Homo sapiens OX=9606 GN=PLOD3 PE=1 SV=1 | O60568 | PLOD3 | 85 kDa | 0 | 1 | 1 | 2 |
| Plasminogen activator inhibitor 1 RNA-binding protein OS=Homo sapiens OX=9606 GN=SERBP1 PE=1 SV=2 | Q8NC51 (+3) | SERBP1 | 45 kDa | 1 | 1 | 0 | 2 |
| 40S ribosomal protein S6 OS=Homo sapiens OX=9606 GN=RPS6 PE=1 SV=1 | P62753 | RPS6 | 29 kDa | 0 | 0 | 2 | 2 |
| E3 ubiquitin-protein ligase HUWE1 OS=Homo sapiens OX=9606 GN=HUWE1 PE=1 SV=3 | Q7Z6Z7 (+2) | HUWE1 | 482 kDa | 1 | 0 | 1 | 2 |
| Serine/arginine repetitive matrix protein 2 OS=Homo sapiens OX=9606 GN=SRRM2 PE=1 SV=2 | Q9UQ35 | SRRM2 | 300 kDa | 0 | 0 | 2 | 2 |
| Nitric oxide synthase, endothelial OS=Homo sapiens OX=9606 GN=NOS3 PE=1 SV=4 | P29474 | NOS3 | 133 kDa | 0 | 1 | 1 | 2 |
| Calpastatin OS=Homo sapiens OX=9606 GN=CAST PE=1 SV=4 | P20810 (+8) | CAST | 77 kDa | 0 | 2 | 0 | 2 |

| | | | | | | | |
|---|---|---|---|---|---|---|---|
| Protein kinase C alpha type OS=Homo sapiens OX=9606 GN=PRKCA PE=1 SV=4 | P17252 | PRKCA | 77 kDa | 0 | 1 | 1 | 2 |
| ATP-dependent RNA helicase DHX15 OS=Homo sapiens OX=9606 GN=DHX15 PE=1 SV=2 | O43143 | DHX15 | 91 kDa | 0 | 1 | 1 | 2 |
| TRIO and F-actin-binding protein OS=Homo sapiens OX=9606 GN=TRIOBP PE=1 SV=3 | Q9H2D6 (+2) | TRIOBP | 261 kDa | 0 | 2 | 0 | 2 |
| Nuclear protein localization protein 4 homolog OS=Homo sapiens OX=9606 GN=NPLOC4 PE=1 SV=3 | Q8TAT6 (+1) | NPLOC4 | 68 kDa | 0 | 0 | 2 | 2 |
| Isoform 4 of Plectin OS=Homo sapiens OX=9606 GN=PLEC | Q15149-4 | PLEC | 516 kDa | 0 | 0 | 2 | 2 |
| Heat shock 70 kDa protein 1A OS=Homo sapiens OX=9606 GN=HSPA1A PE=1 SV=1 | P0DMV8 (+2) | HSPA1A | 70 kDa | 0 | 0 | 2 | 2 |
| Poly [ADP-ribose] polymerase 1 OS=Homo sapiens OX=9606 GN=PARP1 PE=1 SV=4 | P09874 | PARP1 | 113 kDa | 0 | 2 | 0 | 2 |
| Glutaredoxin-3 OS=Homo sapiens OX=9606 GN=GLRX3 PE=1 SV=2 | O76003 | GLRX3 | 37 kDa | 0 | 0 | 2 | 2 |
| Isoform 2 of UDP-glucose:glycoprotein glucosyltransferase 1 OS=Homo sapiens OX=9606 GN=UGGT1 | Q9NYU2-2 | UGGT1 | 175 kDa | 2 | 0 | 0 | 2 |
| Dynamin-binding protein OS=Homo sapiens OX=9606 GN=DNMBP PE=1 SV=1 | Q6XZF7 | DNMBP | 177 kDa | 0 | 2 | 0 | 2 |
| Enhancer of mRNA-decapping protein 4 OS=Homo sapiens OX=9606 GN=EDC4 PE=1 SV=1 | Q6P2E9 (+1) | EDC4 | 152 kDa | 2 | 0 | 0 | 2 |
| Alkyldihydroxyacetonephosphate synthase, peroxisomal OS=Homo sapiens OX=9606 GN=AGPS PE=1 SV=1 | O00116 | AGPS | 73 kDa | 0 | 0 | 2 | 2 |
| 1,4-alpha-glucan-branching enzyme OS=Homo sapiens OX=9606 GN=GBE1 PE=1 SV=3 | Q04446 | GBE1 | 80 kDa | 0 | 0 | 2 | 2 |
| 40S ribosomal protein S18 OS=Homo sapiens OX=9606 GN=RPS18 PE=1 SV=3 | P62269 | RPS18 | 18 kDa | 0 | 0 | 2 | 2 |
| cAMP-dependent protein kinase type II-alpha regulatory subunit OS=Homo sapiens OX=9606 GN=PRKAR2A PE=1 SV=2 | P13861 (+1) | PRKAR2A | 46 kDa | 0 | 0 | 2 | 2 |
| Vacuolar protein sorting-associated protein 52 homolog OS=Homo sapiens OX=9606 GN=VPS52 PE=1 SV=1 | Q8N1B4 | VPS52 | 82 kDa | 0 | 0 | 2 | 2 |

| | | | | | | | |
|---|---|---|---|---|---|---|---|
| Heat shock 70 kDa protein 4 OS=Homo sapiens OX=9606 GN=HSPA4 PE=1 SV=4 | P34932 | HSPA4 | 94 kDa | 0 | 1 | 0 | 1 |
| RNA-binding protein 39 OS=Homo sapiens OX=9606 GN=RBM39 PE=1 SV=2 | Q14498 (+1) | RBM39 | 59 kDa | 0 | 1 | 0 | 1 |
| Actin-related protein 3 OS=Homo sapiens OX=9606 GN=ACTR3 PE=1 SV=3 | P61158 | ACTR3 | 47 kDa | 1 | 0 | 0 | 1 |
| Coatomer subunit beta OS=Homo sapiens OX=9606 GN=COPB1 PE=1 SV=3 | P53618 | COPB1 | 107 kDa | 0 | 1 | 0 | 1 |
| Protein transport protein Sec31A OS=Homo sapiens OX=9606 GN=SEC31A PE=1 SV=3 | O94979 (+7) | SEC31A | 133 kDa | 0 | 1 | 0 | 1 |
| Heterochromatin protein 1-binding protein 3 OS=Homo sapiens OX=9606 GN=HP1BP3 PE=1 SV=1 | Q5SSJ5 (+1) | HP1BP3 | 61 kDa | 1 | 0 | 0 | 1 |
| Nucleolar protein 58 OS=Homo sapiens OX=9606 GN=NOP58 PE=1 SV=1 | Q9Y2X3 | NOP58 | 60 kDa | 0 | 0 | 1 | 1 |
| Serine/arginine-rich splicing factor 7 OS=Homo sapiens OX=9606 GN=SRSF7 PE=1 SV=1 | Q16629 (+3) | SRSF7 | 27 kDa | 0 | 0 | 1 | 1 |
| Filamin-binding LIM protein 1 OS=Homo sapiens OX=9606 GN=FBLIM1 PE=1 SV=2 | Q8WUP2 (+1) | FBLIM1 | 41 kDa | 1 | 0 | 0 | 1 |
| Nuclear pore complex protein Nup153 OS=Homo sapiens OX=9606 GN=NUP153 PE=1 SV=2 | P49790 (+2) | NUP153 | 154 kDa | 0 | 0 | 1 | 1 |
| Inverted formin-2 OS=Homo sapiens OX=9606 GN=INF2 PE=1 SV=2 | Q27J81 (+1) | INF2 | 136 kDa | 1 | 0 | 0 | 1 |
| Aminopeptidase B OS=Homo sapiens OX=9606 GN=RNPEP PE=1 SV=2 | Q9H4A4 | RNPEP | 73 kDa | 0 | 1 | 0 | 1 |
| Aldehyde dehydrogenase, mitochondrial OS=Homo sapiens OX=9606 GN=ALDH2 PE=1 SV=2 | P05091 | ALDH2 | 56 kDa | 1 | 0 | 0 | 1 |
| DNA-directed RNA polymerase II subunit RPB1 OS=Homo sapiens OX=9606 GN=POLR2A PE=1 SV=2 | P24928 | POLR2A | 217 kDa | 0 | 0 | 1 | 1 |
| Heat shock protein 105 kDa OS=Homo sapiens OX=9606 GN=HSPH1 PE=1 SV=1 | Q92598 (+3) | HSPH1 | 97 kDa | 0 | 0 | 0 | 0 |
| Protein SET OS=Homo sapiens OX=9606 GN=SET PE=1 SV=3 | Q01105 (+3) | SET | 33 kDa | 0 | 0 | 0 | 0 |
| Nucleoprotein TPR OS=Homo sapiens OX=9606 GN=TPR PE=1 SV=3 | P12270 | TPR | 267 kDa | 0 | 0 | 0 | 0 |

| | | | | | | | |
|---|---|---|---|---|---|---|---|
| Golgi apparatus protein 1 OS=Homo sapiens OX=9606 GN=GLG1 PE=1 SV=2 | Q92896 (+2) | GLG1 | 135 kDa | 0 | 0 | 0 | 0 |
| 60S ribosomal protein L10a OS=Homo sapiens OX=9606 GN=RPL10A PE=1 SV=2 | P62906 | RPL10A | 25 kDa | 0 | 0 | 0 | 0 |
| Poly(rC)-binding protein 2 OS=Homo sapiens OX=9606 GN=PCBP2 PE=1 SV=1 | Q15366 (+3) | PCBP2 | 39 kDa | 0 | 0 | 0 | 0 |
| Cytochrome b-c1 complex subunit 1, mitochondrial OS=Homo sapiens OX=9606 GN=UQCRC1 PE=1 SV=3 | P31930 | UQCRC1 | 53 kDa | 0 | 0 | 0 | 0 |
| Heterogeneous nuclear ribonucleoprotein D-like OS=Homo sapiens OX=9606 GN=HNRNPDL PE=1 SV=3 | O14979 (+2) | HNRNPDL | 46 kDa | 0 | 0 | 0 | 0 |
| 60S ribosomal protein L14 OS=Homo sapiens OX=9606 GN=RPL14 PE=1 SV=4 | P50914 | RPL14 | 23 kDa | 0 | 0 | 0 | 0 |
| 40S ribosomal protein S5 OS=Homo sapiens OX=9606 GN=RPS5 PE=1 SV=4 | P46782 | RPS5 | 23 kDa | 0 | 0 | 0 | 0 |
| AP-2 complex subunit alpha-2 OS=Homo sapiens OX=9606 GN=AP2A2 PE=1 SV=2 | O94973 (+2) | AP2A2 | 104 kDa | 0 | 0 | 0 | 0 |
| Acidic leucine-rich nuclear phosphoprotein 32 family member A OS=Homo sapiens OX=9606 GN=ANP32A PE=1 SV=1 | P39687 | ANP32A | 29 kDa | 0 | 0 | 0 | 0 |
| Tyrosine-protein phosphatase non-receptor type 12 OS=Homo sapiens OX=9606 GN=PTPN12 PE=1 SV=3 | Q05209 | PTPN12 | 88 kDa | 0 | 0 | 0 | 0 |
| Plexin-D1 OS=Homo sapiens OX=9606 GN=PLXND1 PE=1 SV=3 | Q9Y4D7 (+1) | PLXND1 | 212 kDa | 0 | 0 | 0 | 0 |
| Cleavage and polyadenylation specificity factor subunit 6 OS=Homo sapiens OX=9606 GN=CPSF6 PE=1 SV=2 | Q16630 (+2) | CPSF6 | 59 kDa | 0 | 0 | 0 | 0 |
| CD44 antigen OS=Homo sapiens OX=9606 GN=CD44 PE=1 SV=3 | P16070 (+14) | CD44 | 82 kDa | 0 | 0 | 0 | 0 |
| Eukaryotic translation initiation factor 2 subunit 1 OS=Homo sapiens OX=9606 GN=EIF2S1 PE=1 SV=3 | P05198 | EIF2S1 | 36 kDa | 0 | 0 | 0 | 0 |
| CCHC-type zinc finger nucleic acid binding protein OS=Homo sapiens OX=9606 GN=CNBP PE=1 SV=1 | P62633 (+5) | CNBP | 19 kDa | 0 | 0 | 0 | 0 |
| Cathepsin B OS=Homo sapiens OX=9606 GN=CTSB PE=1 SV=3 | P07858 | CTSB | 38 kDa | 0 | 0 | 0 | 0 |

**Supplementary table 2. Novel potential RNA-binding proteins in HUVEC.**

| Protein | Gene name |
|---|---|
| Basement membrane-specific heparan sulfate proteoglycan core protein | HSPG2 |
| Integrin beta-4 | ITGB4 |
| N-acetylglucosamine-6-sulfatase | GNS |
| Thrombospondin-1 | THBS1 |
| Nidogen-1 | NID1 |
| Endoglin | ENG |
| Nitric oxide synthase, endothelial | NOS3 |
| Abl interactor 1 | ABI1 |
| Ras-interacting protein 1 | RASIP1 |
| EH domain-containing protein 2 | EHD2 |
| Multimerin-2 | MMRN2 |
| Protein kinase C alpha type | PRKCA |
| Fermitin family homolog 3 | FERMT3 |
| Toll-interacting protein | TOLLIP |
| TRIO and F-actin-binding protein | TRIOBP |
| SH3 and multiple ankyrin repeat domains protein 3 | SHANK3 |
| Rho-related GTP-binding protein RhoC | RHOC |
| Vacuolar protein sorting-associated protein 52 homolog | VPS52 |
| Filamin-binding LIM protein 1 | FBLIM1 |
| Aminopeptidase N | ANPEP |

**Supplementary table 3. GO Biological functions of novel potential RNA-binding proteins in HUVEC based on STRING analysis.**

| #term ID | Term description | False Discovery Rate |
|---|---|---|
| GO:0009653 | Anatomical structure morphogenesis | 9.26e-07 |
| GO:0048646 | Anatomical structure formation involved in morphogenesis | 2.68e-06 |
| GO:0001525 | Angiogenesis | 3.36e-06 |
| GO:0035295 | Tube development | 1.60e-05 |
| GO:0022603 | Regulation of anatomical structure morphogenesis | 1.63e-05 |
| GO:0035239 | Tube morphogenesis | 1.71e-05 |
| GO:0048856 | Anatomical structure development | 8.76e-05 |
| GO:0009888 | Tissue development | 0.00027 |
| GO:0050793 | Regulation of developmental process | 0.00097 |
| GO:0048729 | Tissue morphogenesis | 0.0010 |
| GO:0048731 | System development | 0.0016 |
| GO:0043535 | Regulation of blood vessel endothelial cell migration | 0.0020 |
| GO:0007155 | Cell adhesion | 0.0022 |
| GO:0051094 | Positive regulation of developmental process | 0.0022 |
| GO:0045765 | Regulation of angiogenesis | 0.0067 |
| GO:0001936 | Regulation of endothelial cell proliferation | 0.0078 |
| GO:0001937 | Negative regulation of endothelial cell proliferation | 0.0086 |
| GO:0030334 | Regulation of cell migration | 0.0145 |
| GO:0043536 | Positive regulation of blood vessel endothelial cell migration | 0.0145 |
| GO:0045766 | Positive regulation of angiogenesis | 0.0145 |
| GO:0044087 | Regulation of cellular component biogenesis | 0.0166 |
| GO:1903588 | Negative regulation of blood vessel endothelial cell proliferation involved in sprouting | 0.0177 |
| GO:0009790 | Embryo development | 0.0239 |
| GO:0060429 | Epithelium development | 0.0262 |
| GO:0030155 | Regulation of cell adhesion | 0.0411 |
